# Lymphatics constitute a novel component of the intestinal stem cell niche

**DOI:** 10.1101/2022.01.28.478205

**Authors:** Norihiro Goto, Shinya Imada, Vikram Deshpande, Ömer H. Yilmaz

**Author notes:** Correspondence (N.G.), (Ö.H.Y.).

## Abstract

Intestinal stem cells (ISCs) depend on niche factors for their proper function. However, the source of these ISC niche factors and how they support ISCs remain controversial due to their redundant expression patterns. Here, we report that the maintenance of ISCs depends on both lymphatic endothelial cells (LECs) and Rspo3+Grem1+ fibroblasts (RGFs). We found that LECs are surrounded by RGFs and located in close proximity to Lgr5+ ISCs. RSPO3 production is restricted to LECs and RGFs and they can partially compensate for each other; however, RSPO3 loss in both of LECs and RGFs drastically compromises ISC numbers, villi length, and repair after irradiation-induced injury. Mechanistically, irradiation-induced damage expands LEC and RGF numbers and enhances the latter’s generation of RSPO3 through IL-1 receptor activation. We propose that LECs represent a novel component of the ISC niche, which together with RGFs, provide essential RSPO3 to sustain ISCs in homeostasis and regeneration.

## INTRODUCTION

Lgr5+ intestinal stem cells (ISCs) self-renew and give rise to the various specialized cell types of the intestinal epithelium (Gehart and Clevers, 2019). ISCs reside at crypt bases and depend on niche factors such as Wnt, R-spondins (RSPO), BMP inhibitors (BMPi), and EGF for maintaining the stem cell state (Sato et al., 2009). The essentiality of these factors, for example, in supporting ISC function has been defined by their need to support intestinal organoid cultures. These ISC factors are produced by many of their neighboring cell types including epithelial and mucosal stromal cell types (Degirmenci et al., 2018; Greicius et al., 2018; McCarthy et al., 2020b; Shoshkes-Carmel et al., 2018). Epithelial Paneth and deep secretory cells adjacent to Lgr5+ ISCs in the small intestine and colon, respectively, produce Wnt3 and EGF to support Lgr5+ ISCs (Sasaki et al., 2016; Sato et al., 2011; Yilmaz et al., 2012); however, there are redundant sources for Wnt, including stromal cells -where co-culture of stromal cells with Wnt3 deleted organoids permits organoid propagation (Farin et al., 2016; Farin et al., 2012). Finally, the absence of RSPO and BMPi production in epithelial cells further accentuates the role that stromal cells play as key constituents of the ISC niche (Greicius et al., 2018; Kosinski et al., 2007; Ogasawara et al., 2018).

In an attempt to define the stromal ISC niche, several mouse models and strategies to genetically manipulate or sort populations of stromal cells by cell surface markers have been developed and employed (Degirmenci et al., 2018; Greicius et al., 2018; Harnack et al., 2019; McCarthy et al., 2020a; McCarthy et al., 2020b; Shoshkes-Carmel et al., 2018; Stzepourginski et al., 2017). However, the source of these ISC niche factors and how they support ISCs in vivo still remain controversial. First, direct visualization of these factors with precise cellular localization is technically challenging for low expressing genes or secreted factors. These approaches include in situ hybridization or immunohistochemical approaches, which also fail to provide functional insights. Second, genetically engineered mouse models used in these studies (e.g., *Pdgfra-Cre, Foxl1-Cre, Gli1-CreERT2* mice) do not mark cell populations that produce specific ISC niche factors and many ISC niche factors are expressed by more than one cell type, making it difficult to ascribe in vivo loss of function phenotypes to specific cell types (Degirmenci et al., 2018; Greicius et al., 2018; McCarthy et al., 2020a; Shoshkes-Carmel et al., 2018). For example, telocytes, marked by Foxl1+ cells (Shoshkes-Carmel et al., 2018), Gli1+ cells (Degirmenci et al., 2018) or Pdgfra^high^ cells (McCarthy et al., 2020b) in the stroma, have been proposed to serve as a major source of Wnt in the intestinal stroma. Loss of active Wnt in telocytes reduced crypt cell proliferation in the intestine, indicating that these cells are essential in providing Wnt to intestinal stem and progenitor cells (Degirmenci et al., 2018; Shoshkes-Carmel et al., 2018). While *Porcn* deletion in Foxl1+ cells (i.e. telocytes) led to rapid crypt collapse in 3 days both in the small intestine and colon (Shoshkes-Carmel et al., 2018), *Wls* deletion in Gli1+ cells (i.e. telocytes) had minimal effect in the small intestine and necessitated 2-3 weeks to cause crypt attrition in the colon (Degirmenci et al., 2018). These distinct phenotypes likely reflect that Gli1 and Foxl1 positive stromal cells mark partially overlapping populations of cells as revealed by single cell analysis (Degirmenci et al., 2018; Kinchen et al., 2018; McCarthy et al., 2020b; Shoshkes-Carmel et al., 2018), highlighting the challenges of precisely manipulating growth factors using single promoter driven Cre recombinase models. Trophocytes, defined as CD81+Pdgfra^low^ expressing fibroblast, are enriched for Grem1+ cells and support organoid growth through the production of BMPi (namely Grem1) and RSPO3 (McCarthy et al., 2020b). Although diphtheria toxin mediated cellular ablation of Grem1+ cells compromises ISC function, it is unclear whether ISC loss results from the lack of Grem1 production, loss of other growth factors such as RSPO in those cells, or through non-specific causes as Grem1 is also expressed by other stromal fibroblasts (non-trophocytes) and the musclaris propria (i.e. loss of intestinal integrity). Thus, the development of additional tools to investigate and visualize specific stromal cell populations that correspond to specific ISC niche factors in vivo may help to clarify their in vivo role in fostering ISCs.

RSPOs boost Wnt signaling by functioning as Lgr5 ligands (de Lau et al., 2014). RSPOs drive self-renewal of Lgr5+ ISCs *in vivo* (Yan et al., 2017) and are indispensable trophic factors for intestinal organoid growth *in vitro* (Sato et al., 2009). RSPOs are encoded by four paralogous genes: *Rspo1-Rspo4* (Greicius et al., 2018). RSPO3 is the dominant R-spondin in the mammalian intestine and is significantly more potent that RSPO1 in enhancing Wnt activation and supporting intestinal organoid growth (Greicius et al., 2018). Previous studies have proposed that Rspo3 is expressed by subepithelial myofibroblasts (Greicius et al., 2018; Stzepourginski et al., 2017), a subset that was later identified as telocytes (Shoshkes-Carmel et al., 2018), by Pdgfra^+^ fibroblasts (Greicius et al., 2018; McCarthy et al., 2020b), or by lymphatic endothelial cells (LECs) (Ogasawara et al., 2018); however, questions remain regarding which of these cell types is the major RSPO3 source for ISCs as previous studies have reported that Rspo3 deletion has minimal impact on intestinal homeostasis and integrity (Greicius et al., 2018; Harnack et al., 2019). In these studies, RSPO3 was either ablated during embryonic development or in a subset of RSPO3 producing cells, using *Pdgfra-Cre; Rspo3 f/f* (Greicius et al., 2018) or *Myh11-CreERT2; Rspo3 f/f* genetically engineered mice, respectively (Harnack et al., 2019). The first model disrupts *Rspo3* with a constitutive Cre in a broad population of Pdgfra+ intestinal stromal cells and may be confounded by compensation from other RSPO family members given early embryonic deletion, and the second model disrupts *Rspo3* in Myh11+ myofibroblasts, a population that does not show robust *Rspo3* expression by scRNA-seq (Brugger et al., 2020; McCarthy et al., 2020b). While in both of these models Rspo3 loss either has mild effects of Lgr5+ ISC numbers and intestinal integrity in homeostasis (Greicius et al., 2018; Harnack et al., 2019), systemic overexpression of RSPO receptors that sequester in vivo RSPO proteins causes notable crypt degeneration (Yan et al., 2017), raising the possibility that *Rspo3* has not been completely excised from the mucosal ISC niche in these in vivo loss of function models. Finally, it has been previously noted that LECs are RSPO3+ but whether or how they support ISCs has not been explored (Ogasawara et al., 2018).

To address these issues, we employed multiple novel reporter mice to directly visualize and sort ISC niche factor-producing cells as well as to perform loss of function studies. We find that LECs represent a novel niche component for ISCs, which together with RSPO3+GREM1+ fibroblasts, serve as the major in vivo RSPO3 source for ISCs in homeostasis and injury-mediated regeneration.

## RESULTS

### *Rspo3*+ LECs reside in close proximity to *Lgr5*+ ISCs

To visualize and characterize ISC niche factor-producing cells, we first focused on RSPO3 and generated *Rspo3-GFP* BAC transgenic mice. Immunostaining for *Rspo3*-GFP highlighted endothelial cell-like structure in the small intestine and colon (Figure 1A). This distribution is consistent with the *Rspo3* mRNA expression pattern detected by in situ hybridization (ISH) (Figure S1A). To distinguish between LECs and vascular endothelial cells, we performed immunofluorescence for LEC-specific marker LYVE-1 and *Rspo3-GFP* and found that Rspo3 is expressed by LYVE-1+ LECs (Figure 1A), which is consistent with a prior study (Ogasawara et al., 2018).

**Figure 1.**
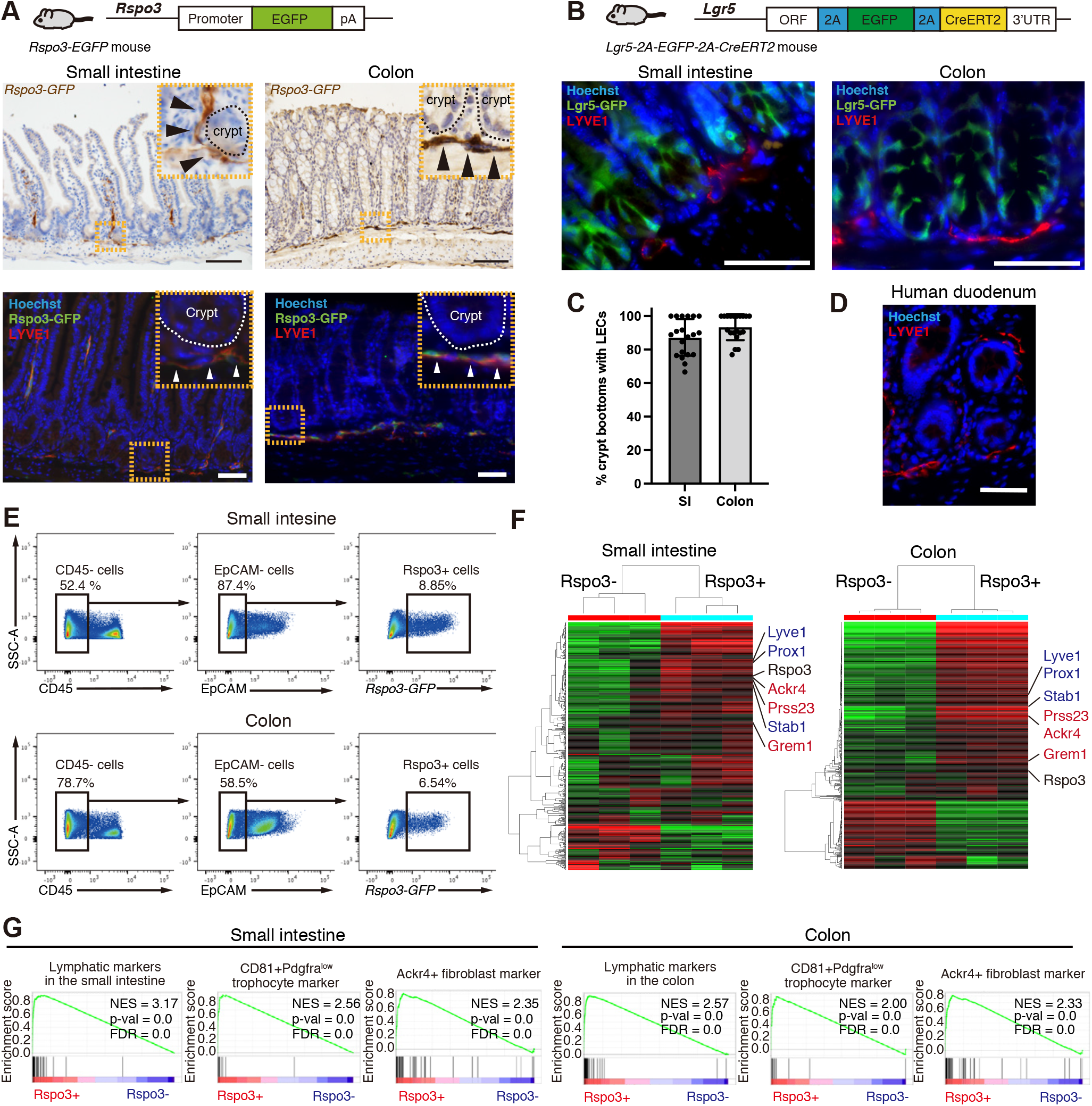
*Rspo3*+ LECs reside in close proximity to *Lgr5*+ ISCs. (A) Schematic of Rspo3-GFP mice (top). Rspo3-GFP expression by immunohistochemistry in the small and colon (middle). Black arrowheads indicate endothelial structure in close proximity to the crypt. Immunofluorescence (IF) shows Rspo3 is expressed by LYVE1+ Lymphatic endothelial cells (LECs) (bottom). White arrowheads indicate the co-expression of Rspo3-GFP and LYVE1. The image represents one of 6 biological replicates. (B) Schematic (top) of new *Lgr5-2A-GFP-2A-CreERT2* mice. IF for Lgr5-GFP and LYVE1 in the small intestine and colon shows the presence of LYVE1+ LECs residing in close proximity to Lgr5+ ISCs. The image represents one of 6 biological replicates. (C) The percentage of the crypts with adjacent LYVE1+ LECs in the small intestine and the colon. n = 20 high-power fields from 6 mice per group. (D) IF for LYVE1 in human duodenum. The image represents one of 10 biological replicates. (E) CD45^−^/EpCAM^−^/Rspo3-GFP^+^ cells from the small intestinal and colonic stromal cells of Rspo3-GFP mice by flow cytometry. The plots represent one of >10 biological replicates. (F) The heatmap of RNA-seq of sorted CD45^−^/EpCAM^−^/Rspo3-GFP^−^ and CD45^−^/EpCAM^−^/Rspo3-GFP^+^ cells in the small intestine and colon. n = 3 mice per group. (G) Gene Set Enrichment Analysis (GSEA) of lymphatic markers, CD81^+^Pdgfra^low^ trophocyte markers, and Ackr4+ fibroblast markers. FDR, false-discovery rate; NES, normalized enrichment score. Scale bar, 50 μm. Data are mean ± SD.

Given the role that RSPO3 plays in driving intestinal stemness, we next investigated the localization of LECs in relation to *Lgr5*+ ISCs. Because conventional *Lgr5-EGFP-IRES-CreERT2* mice (Barker et al., 2007) have issues of variegated reporter expression (Schuijers et al., 2014), we generated *Lgr5-2A-EGFP-2A-CreERT2* mice (Figure S1B). In this new *Lgr5* reporter, *Lgr5-GFP* is expressed in all small intestinal and colonic crypts (Figure S1B). We also confirmed that these *Lgr5-GFP*+ cells self-renew and differentiate for the long-term in the small intestine and colon by performing lineage tracing using *Lgr5-2A-EGFP-2A-CreERT2; Rosa-LSL-tdTomato* mice (Figure S1C). Immunofluorescence and confocal microscopy confirmed the presence of LYVE1+ LECs residing in close proximity to small intestinal and colonic Lgr5+ ISCs (Figure 1B and movie S1) in mice (Figure 1C) and human samples (Figure 1D). In the colon, LYVE1+ LECs are located near the crypt base, whereas in the small intestine, LYVE1+ LECs are located both at the crypt base and in the villious cores (also known as villous lacterals) (Cifarelli and Eichmann, 2019) (Figure 1A). These lacterals not only reside far from the epithelium but soluble Rspo3 likely exerts minimal effects on the *Lgr5* non-expressing differentiated villous epithelium.

Next, we performed RNA-seq on sorted *Rspo3-GFP*- and *Rspo3-GFP*+ cells from EpCAM-CD45-stromal cells in the small intestine and in the colon (Figures 1E and 1F). Differential expression analysis and Gene Set Enrichment Analysis (GSEA) showed enrichment of LEC marker genes (e.g. *Lyve1*, *Prox1*, *Stab1*) and *Grem1+* fibroblast (e.g., CD81^+^Pdgfra^low^ trophocyte (McCarthy et al., 2020b) and Ackr4+ fibroblast (Thomson et al., 2018)) marker genes (*Ackr4*, *Grem1*, *Prss23)* (McCarthy et al., 2020b; Thomson et al., 2018) (Figure 1F, G), demonstrating heterogeneity of Rspo3+ cells. Reanalysis of single-cell RNA sequencing (scRNA-seq) data of the small intestinal (McCarthy et al., 2020b) and colonic (Kinchen et al., 2018) stroma also showed that *Rspo3* is not only expressed by LECs but also by a subset of *Pdgfra^low^* fibroblasts that co-express *Grem1* (Figures S2A and S2B). We defined these Rspo3+Grem1+ fibroblasts as “RG-fibroblasts (RGFs)”.

### LECs and RGFs are the major source of mucosal Rspo3

To resolve the heterogeneity of Rspo3+ stromal cells, we performed scRNA-seq on sorted small intestinal *Rspo3-GFP*+ stromal cells (Figure 2A). We detected two dominant clusters: a large cluster of *Cd31*+ cells (46%; 1,739/3,774) that express LEC markers (LEC cluster) and another cluster of *Cd31*-cells (52.5%; 1,983/3,774) that express *Grem1* (RGF cluster) (Figure 2B). In addition, we also detect a very small cluster of telocytes (1.3%; 51/3774) that highly express *Pdgfra* as well as *Foxl1*, *Bmp3* and *Bmp5* (Figure 2B), as previously reported (McCarthy et al., 2020b; Shoshkes-Carmel et al., 2018). Given that the telocyte cluster is quite small, telocytes likely account for a small proportion of Rspo3 production, in contrast to LECs and RGFs. There are 4 subclusters in LECs and 5 subclusters in RGFs (Figure 2A); all the subclusters are similar in terms of the production of ISC niche factors (Figure 2D). In RGFs, one cluster (RGF 2) is Cd81−Adamdec1+ while other four clusters (RGF 1-1, 1-2, 1-3, 1-4) are Cd81+Adamdec1− (Figure 2C). The former corresponds to a subset of CD81−Pdgfra^low^ fibroblasts, while the latter corresponds to a subset of CD81+Pdgfra^low^ trophocytes. We confirmed the distribution of Rspo3+Grem1+ cells (RGFs) in both the Cd81−Pdgfra^low^ fibroblast and Cd81+Pdgfra^low^ trophocyte clusters in the reanalysis of scRNA-seq data of small intestinal Pdgfra+ cells (McCarthy et al., 2020b) (Figure S3A).

**Figure 2.**
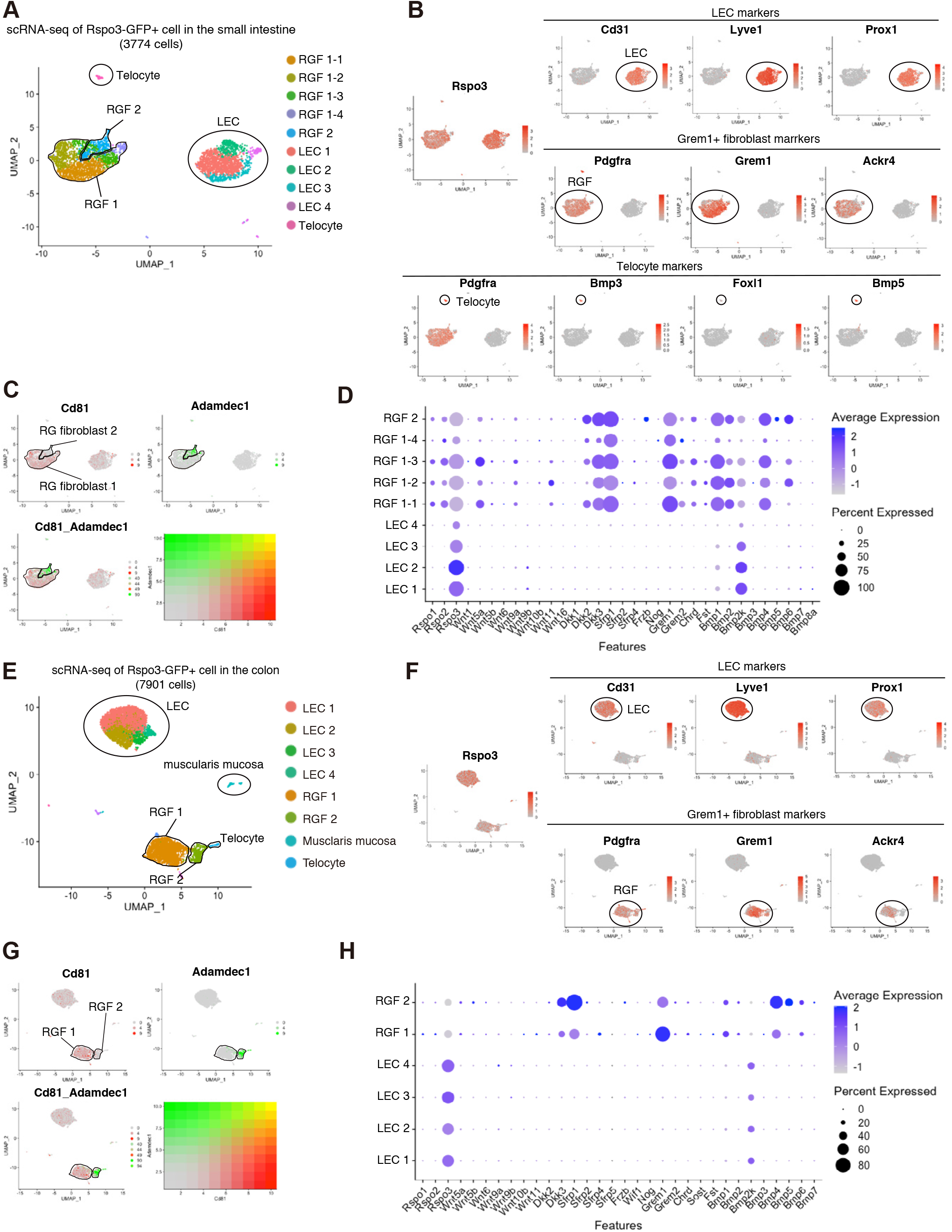
scRNA-seq of Rspo3+ cells in the small intestine and colon. **(A)** Uniform manifold approximation and projection (UMAP) of scRNA-seq of small intetstinal Rspo3− GFP+ cells (3,774 cells, n = 2 mice). **(B)** Relative expression of *Rspo3*, LEC marker genes, and Grem1+ fibroblast marker genes, and telocyte marker genes onto the UMAP plot. **(C)** Relative expression of *Cd81* and *Adamdec1* on *the* UMAP plot, showing that RGF 1 clusters are CD81+Adamdec1− whereas RGF 2 cluster is Cd81−Adamdec1+. **(D)** Dot plots of expression patterns of intestinal stem cell niche factors in LEC and RGF subclusters. **(E)**UMAP of scRNA-seq of colonic Rspo3-GFP+ cells (7,901 cells, n = 2 mice). **(F)**Relative expression of *Rspo3*, LEC marker genes, and Grem1+ fibroblast marker genes onto the UMAP plot. **(G)**Relative expression of Cd81 and Adamdec1 on the UMAP plot, showing that RGF 1 cluster is CD81+Adamdec1− whereas RGF 2 cluster is Cd81−Adamdec1+. **(H)** Dot plots of expression patterns of intestinal stem cell niche factors in LEC and RGF subclusters.

We also performed scRNA-seq in colonic *Rspo3-GFP*+ stromal cells. As in the small intestine, we detected two dominant clusters: LEC and RGF clusters (Figure 2E and 2F). Besides these main cluster, there were small clusters of telocytes (0.89%; 71/7,901) and muscularis mucosa cells (1.67%; 132/7,901), the latter of which also express both Grem1 and Rspo3 (Figures 2E and 2F). Since these clusters are small, the major sources of mucosal Rspo3 are LECs and RGFs in the colon akin to the small intestine. There are 4 subclusters in LECs and 2 subclusters in RGFs (Figure 2E); all the subclusters are similar in terms of the production of ISC niche factors (Figure 2H). As noted in the small intestine, RGFs can be subdivided into Cd81+Adamdec1− (RGF 1) and Cd81−Adamdec1+ (RGF 2) subclusters (Figure 2G), which correspond to a subset of CD81−Pdgfra^low^ fibroblasts and a subset of CD81+Pdgfra^low^ trophocytes, respectively (Figure S3B) (Brugger et al., 2020). We also reanalyzed the scRNA-seq of human colon stromal cells (Kinchen et al., 2018). RSPO3+GREM1+ cells (namely RGFs) were again identified in a subset of PDGFRA^low^ cells (Figure S3C) similar to the mouse intestine.

We next separately sort LECs and RGFs (Figure 3A) by utilizing *Rspo3-GFP* and the cell surface marker CD31 that is expressed by LEC but not by RGFs (Figures 2B and 2F). RNA-seq on CD31-*Rspo3-GFP-*, CD31+*Rspo3-GFP-*, CD31-*Rspo3-GFP+*, and CD31+*Rspo3-GFP+* cells confirmed that *Grem1* is essentially expressed by CD31-*Rspo3-GFP+* cells (Figure 3B). GSEA and single sample GSEA (ssGSEA) (Barbie et al., 2009) revealed that CD31-*Rspo3-GFP*+ cells are enriched in RGF markers shared by CD81+Pdgfra^low^ cell (McCarthy et al., 2020b) or Ackr4+ cells (Thomson et al., 2018); CD31+*Rspo3-GFP*+ cells in LEC markers (Kalucka et al., 2020); and CD31+*Rspo3-GFP*-cells in vascular endothelial cell markers (Kalucka et al., 2020) (Figures S4A and S4B), illustrating that we can isolate LECs and RGFs by sorting CD31+*Rspo3-GFP*+ and CD31-*Rspo3-GFP*+ cells, respectively. Because bulk RNA-seq allows for the detection of lowly expressed genes, we next sought to assess the expression of ISC niche factors in LECs and RGFs (Figure S4C-I). Of note, LECs express *Wnt2*, a canonical Wnt (Goss et al., 2009) that supports ISCs, and RGFs express *Wnt11, Wnt2b, Wnt5a, Wnt5b*, and *Wnt9a* (Figure S5I), implying their role to support ISCs through Wnt production as well. Further, RGFs also highly express *Igf1* and *Fgf7* (Figure S4D), growth factors that enhance organoid propagation (Fujii et al., 2018).

**Figure 3.**
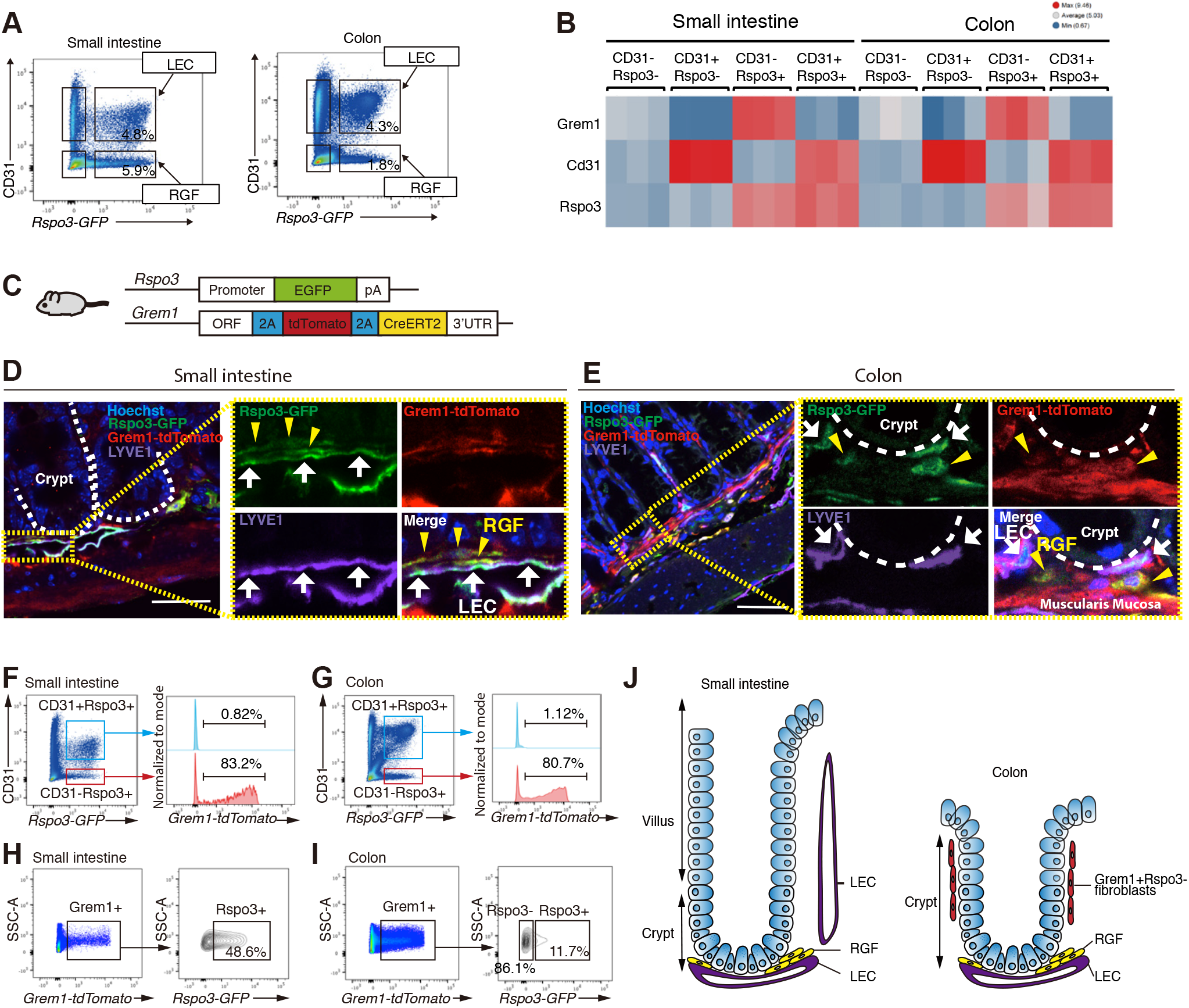
LECs and RGFs are the major source of mucosal Rspo3. (**A**) Flow cytometry of EpCAM^−^CD45^−^ stromal cells using Rspo3-GFP and CD31 in the small intestine and colon. The plots represent one of >10 biological replicates. (**B**) The heatmap of RNA-seq on CD31-*Rspo3-GFP-*, CD31+*Rspo3-GFP-*, CD31-*Rspo3-GFP+*, and CD31+*Rspo3-GFP+* cells in the small intestine and colon. n = 3 mice per group. **(C)** Schematic of *Rspo3-GFP; Grem1-tdTomato* mouse. (**D-E)** Confocal microscopy images of immunofluorescence for Rspo3-GFP, Grem1-tdTomato, and LYVE1 in the small intestine (**D**) and colon (**E**). Yellow arrowheads indicate RGFs. White arrows indicate LEC. The image represents one of 6 biological replicates. **(F-I)** Flow cytometry of EpCAM^−^CD45^−^ stromal cells in the small intestine and colon of *Rspo3-GFP; Grem1-tdTomato* mice. The plots represent one of 3 biological replicates. (**J**) Schematic of Rspo3 and/or Grem1 positive cells in the small intestine and colon. Scale bar, 10 μm (**D**) and 50 μm (**E**).

### RGFs reside close to crypt bottoms and surround lymphatic vasculature

To delineate the location of RGFs, we engineered *Grem1-2A-tdTomato-2A-CreERT2* mice (Figure S5A). *Grem1-tdTomato* is expressed by a subset of lamina propria and submucosal stromal cells, but is also strongly expressed by Desmin+ muscularis propria and muscularis mucosa cells, the latter of which is a thin layer of muscle that separates the mucosal lamina propria from the submucosa (Figure S5B). To visualize RGFs in the lamina propria, we generated *Rspo3-GFP; Grem1-tdTomato* mice (Figure 3C). Confocal microscopy of the small intestine revealed that RGFs are detected in close proximity to LYVE+ LECs near the crypt bottom cells (Figure 3D) and are infrequently detected in the villous core next to lacteal LECs (Figure S5C). These villous RGFs co-express Adamdec1 (Figure S5C), indicating that they correspond to the RGF cluster 2 (CD81−Adamdec1+ subcluster of RGFs) (Figure 2C). In the colon, RGFs are again detected adjacent to LYVE+ LECs both in the lamina propria above the muscularis mucosa and in the submucosal layer below the muscularis mucosa (Figure 3E). Given the close proximity of LECs and RGFs to the crypt base where ISCs resides, these stromal cells likely represent the main of source of Rspo3 and Grem1 for ISCs. Further away from the crypt bottom and consistent with our scRNA-seq analysis (Figure 2E), a small subset of Rspo3+Grem1+ cells are detected in muscularis mucosa (Figure 3E), where they serve as a redundant source of these niche factors. Of note, we also detected many Grem1+Rspo3− cells at the middle-top zone of colonic crypts but not in the small intestine (Figures S5D and 3J).

We also performed flow cytometry of EpCAM^−^CD45^−^ stromal cells in *Grem1-tdTomato; Rspo3-GFP* mice. More than 80% of CD31-*Rspo3-GFP*+ cells express Grem1 both in the small intestine and colon whereas CD31+*Rspo3-GFP*+ cells do not express Grem1 (Figures 3F and 3G). While half of Grem1+ cells express Rspo3 in the small intestine, 86% of Grem1+ cells do not express Rspo3 in the colon (Figures 3H and 3I), which corresponds to Grem1+Rspo3− cells in the middle-top colonic crypts (Figures S5D and 3J). Thus, it is not possible to enrich for RGFs based solely on Grem1 expression. Although the distributions of Rspo3 and/or Grem1 positive cells differs in the differentiated regions of the villi or upper crypts of the small intestine and colon, respectively (Figure 3J), LECs and RGFs consistently reside adjacent to the ISC zone of the crypt base, highlighting the likely critical role that they play in fostering ISCs.

The proximity of RGFs to LECs prompted us to explore possible cross-regulation between these two populations. LECs express *Vegfr2* and *Vegfr3*, whereas their ligands, *Vegfc* and *Vegfd*, are highly expressed by RGFs (Figure S4C). RGFs may, thus, support the growth of LECs through VEGFC and VEGFD production, essential factors required for lymphangiogenesis (Karaman et al., 2018). Further, as we mentioned above, RGFs also highly express *Igf1* and *Fgf7* (Figure S4D), growth factors that not only enhance organoid propagation (*28*) but also aid LEC function in vitro and in vivo (Bjorndahl et al., 2005; Chang et al., 2004; Lokmic, 2016), indicating that they likely play a regulatory role in ISC and LEC biology.

### LECs and RGFs support ISCs in vitro

Next, to ascertain whether Rspo3+ stromal cells support ISCs, we set out to perform heterotypic co-culture experiments with Rspo3+ stromal cells and intestinal epithelial crypts. We adapted a protocol for the culture of dermal lymphatic endothelial cells (Lokmic, 2016), and successfully propagate sorted *Rspo3-GFP*+ cells both in 2-D and 3-D cultures (Figures S5E and S5F). We sorted CD31+*Rspo3-GFP*+ LECs and CD31-*Rspo3-GFP*+ RGFs and then performed heterotypic co-culture experiments with intestinal crypts in the culture media supplemented with Noggin, but not with RSPO (Figure 4A). Organoid formation was confirmed both with LECs and RGFs (Figure 4B). The organoid forming capacity was significantly higher with RGFs than with LECs (Figure 4C), though the latter was still able to aid organoid propagation; this difference between these two populations likely reflects that RGFs expand significantly more in culture than LECs (Figures S5G and S5H). In support of this, we confirmed that Rspo3+ stromal cells support organoid growth in a cell number-dependent manner (Figures S5I-K)

**Figure 4.**
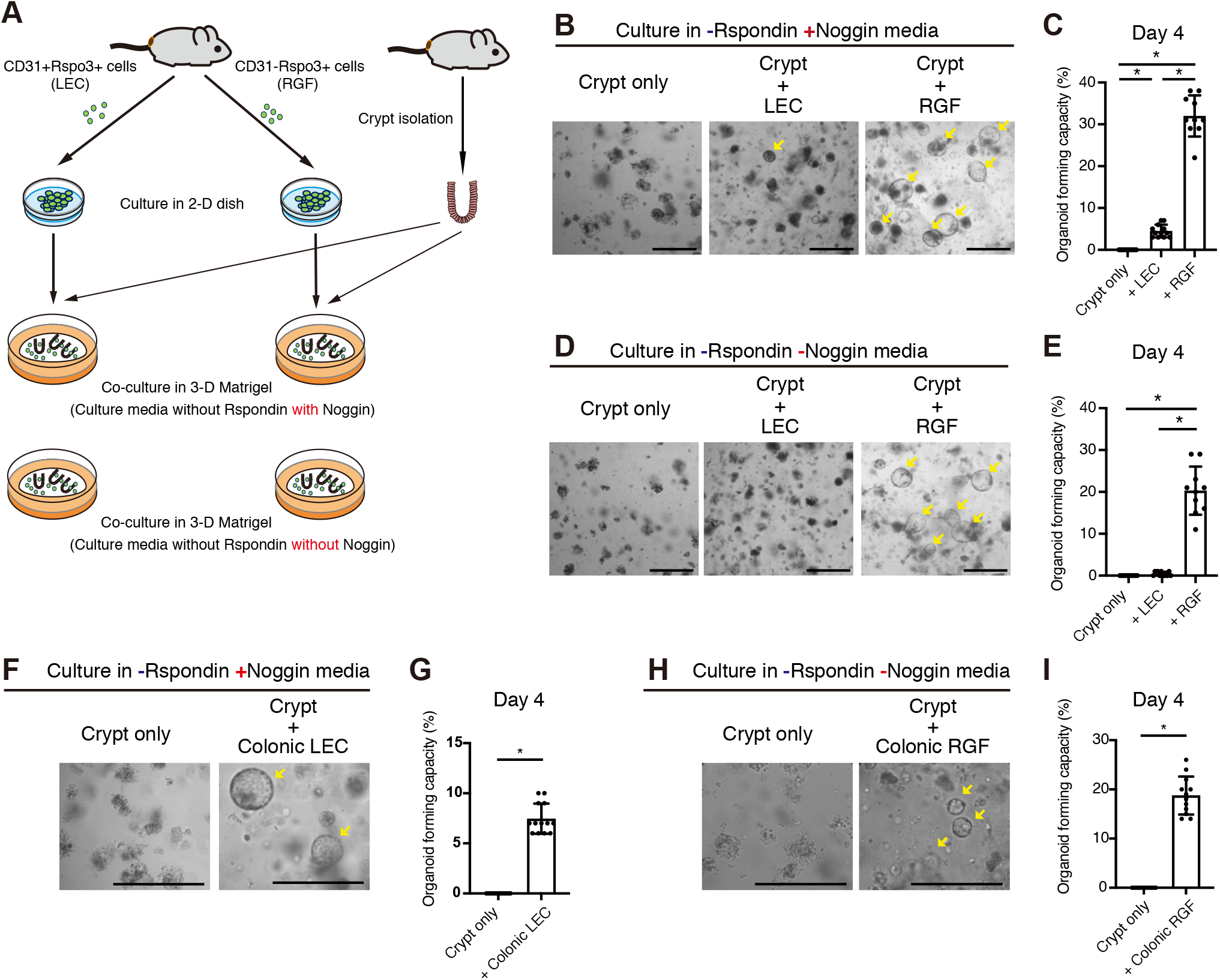
LECs and RGFs support intestinal stem cells in vitro. **(A)** Schematic of heterotypic co-culture. **(B-C)**, Representative images (**B**) and quantification (**C**) of co-culture of small intestinal LECs or RGFs with the crypts in the culture media supplemented with Noggin, but not with RSPO. n = 10 - 12 from 3 mice per group. **(D-E)**, Representative images (**D**) and quantification (**E**) of co-culture of small intestinal LECs or RGFs with the crypts in the culture media without Noggin and RSPO. n = 10 - 12 from 3 mice per group. **(F-G)**, Representative images (**F**) and quantification (**G**) of co-culture of colonic LECs with the crypts in the culture media supplemented with Noggin, but not with RSPO. n = 13 from 3 mice per group. **(H-I)**, Representative images (**H**) and quantification (**l**) of co-culture of colonic RGFs with the crypts in the culture media without Noggin and RSPO. n = 12 from 3 mice per group. One-way analysis of variance (ANOVA) with post-hoc Tukey’s multiple comparison (**C, E**). Unpaired two-tailed t-tests (**G, I**). Data are mean ± SD. *p < 0.05. Scale bar, 20 μm (**B, D, F, H**). Arrows indicate organoid formation.

Because RGFs also produce BMPi Grem1, we deciphered whether RGFs can substitute for Rspo3 and BMPi Noggin. Whereas LECs did not support the organoid formation in culture media that lacks both Rspo3 and Noggin, RGFs robustly sustained organoid growth (Figures 4D and 4E), confirming that RGFs secrete two key niche factors namely Rspo3 and Grem1 to support ISCs. Finally, we also validated that colonic LECs akin to their small intestinal counterparts can support organoid initiation in culture media that lacks Rspo (Figures 4F and 4G) and that colonic RGFs can support organoid initiation in culture media that lacks RSPO and Noggin (Figures 4H and 4I).

### LECs and RGFs support ISCs and post-injury regeneration in vivo

We next sought out to ascertain the in vivo role and sources of Rspo3 in ISC maintenance and repair after injury. We took advantage of our newly engineered *Grem1-CreERT2* driver (Figure S5A) and obtained *Prox1-CreERT2* (a driver specific to lymphatic endothelium (Srinivasan et al., 2007)) and *Rspo3 flox* mice (Neufeld et al., 2012) to ablate Rspo3 in RGFs and LECs, respectively, its primary sources near ISCs. To conditionally disrupt Rspo3 in LECs, we generated *Prox1-CreERT2; Rspo3 f/f* mice, and in RGFs, generated *Grem1-CreERT2; Rspo3 f/f* mice, and in both LECs and RGFs, generated *Grem1-CreERT2; Prox1-CreERT2; Rspo3 f/f* mice (Figures 5A and 5B). Rspo3 loss in either LECs or RGFs diminished numbers of Olfm4+ or Lgr5+ ISCs, but did not show any architectural changes in the small intestine and the colon (Figures 5C-J), indicating compensation of Rspo3 from the unexcised niche population. In support of this notion, when Rspo3 is deleted in both LECs and RGFs using *Grem1-CreERT2; Prox1-CreERT2; Rspo3 f/f* mice, we now observed a dramatic decrease in the numbers of Olfm4+ or Lgr5+ ISCs that is accompanied by shortened crypt-villus units and crypts in the small intestine and the colon, respectively (Figures 5C-J). These phenotypical changes emulate the morphological changes observed in a new model of diphtheria toxin mediated ablation of Lgr5+ ISCs (Tan et al., 2021).

**Figure 5.**
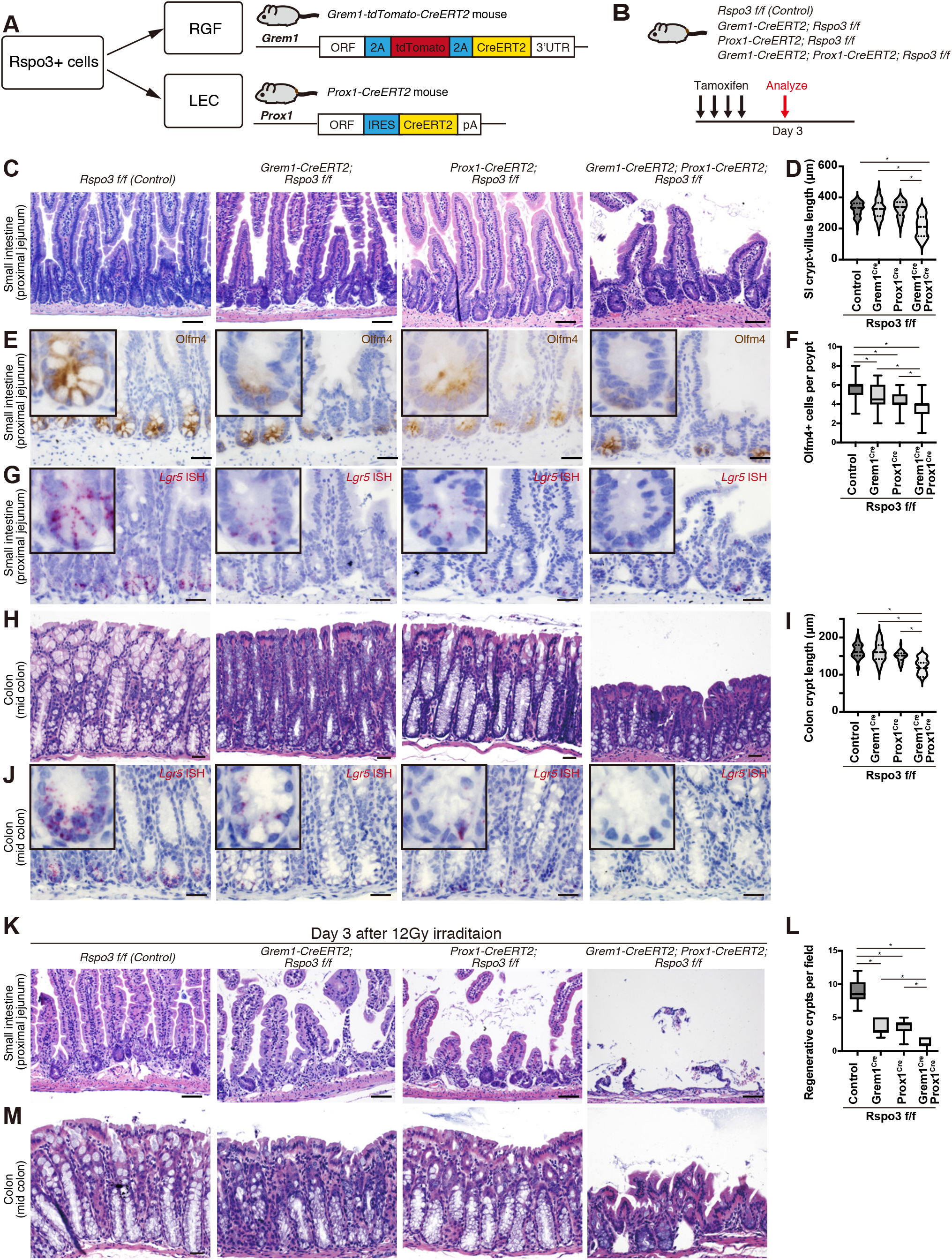
LECs and RGFs support ISCs and post-injury regeneration in vivo. (**A**) Schematic of the Cre mouse models to target Rspo3+ cells. (**B**) Schematic of the mouse models of Rspo3 loss in LECs, RG fibroblasts, or both. **(C-J)**, H&E staining of the small intestine (**C**) and colon (**H**); immunohistochemistry for Olfm4 in the small intestine (**E**); *Lgr5* mRNA expression in the small intestine (**G**) and colon (**J**) by ISH after inducing Rspo3 loss in LECs, RGFs, or both. Quantification of the crypt-villus length in the proximal jejunum of the small intestine (**D**) (n = 100 crypt-villi from 6 mice per group); Olfm4+ cells in the proximal jejunum of the small intestine (**F**) (n = 60 crypts from 6 mice per group); crypt length in the mid colon (**I**) (n = 40 crypts from 6 mice) after inducing Rspo3 loss in LECs, RGFs, or both. (**K-M**), H&E staining of the small intestine (**K**) and colon (**M**) and quantification of regenerative crypts in the proximal jejunum (**L**) (n = 10 fields from 6 mice) day 3 post-irradiation following Rspo3 loss in LECs, RGFs, or both. The images represent one of 6 biological replicates per group (**C**, **E**, **G**, **H**, **J**, **K**, **M**). One-way analysis of variance (ANOVA) with post-hoc Tukey’s multiple comparison (**D**, **F**, **I**, **L**). For box-and-whisker plots (**F**, **L**), data were expressed as box-and-whisker from the minimum to the maximum. *p < 0.05. Scale bar, 20 μm (**C**, **E**, **G**, **H**, **J**, **K**, **M**).

To assess how injury affects compromised Lgr5+ ISC function upon Rspo3 loss in LECs, RGFs or both, we exposed these mice to 12 Gy irradiation after tamoxifen administration. At day 3 post-injury, whereas enlarged/hyperplastic regenerative crypts are seen in the control mice, areas crypt attrition/loss lesions and reduced regenerative crypts were enumerated in the *Grem1-CreERT2; Rspo3 f/f* and *Prox1-CreERT2; Rspo3 f/f* models (Figures 5K and 5L). When Rspo3 is disrupted in both LECs and RGFs (that is, in *Grem1-CreERT2; Prox1-CreERT2; Rspo3 f/f* mice), there is a pronounced increase in ulcerated mucosa that lacks intact crypts (Figures 5K and 5L). Since the colonic epithelium is more resistant to irradiation induced damage as compared to the small intestine (Hua et al., 2017), colonic crypt attrition/loss day 3 post-injury was not detected in control, *Grem1-CreERT2; Rspo3 f/f* or *Prox1-CreERT2; Rspo3 f/f* mice but was only seen when Rspo3 was ablated in both LECs and RGFs (Figure 5M). These results demonstrate that LECs and RGFs play an overlapping role in providing niche Rspo3 role to Lgr5+ ISC in homeostasis and post-injury repair.

### LECs and RGFs expand to facilitate epithelial regeneration after irradiation induced damage

Given the significant role of LEC and RGF derived Rspo3 in ISC function after injury, we next investigated how these niche cells respond to irradiation mediated injury. Interestingly, irradiation expanded lymphatic vascular channels near the crypt based as noted by histologic examination (Figure 5K) and immunofluorescence for LYVE1 (Figure 6A). We independently validated that CD31+*Rspo3− GFP*+ LECs and CD31-*Rspo3-GFP*+ RGFs were expanded by flow cytometric quantification day 3 post-irradiation (Figures 6B and 6C). To better visualize the expanded RGFs, we similarly irradiated *Rspo3-GFP; Grem1-tdTomato* mice and found by confocal microscopy that LECs and RGFs are present in greater quantities near regenerating crypts (Figure 6D), indicating that ISC mediated regeneration is accompanied by stromal niche augmentation in this injury model.

**Figure 6.**
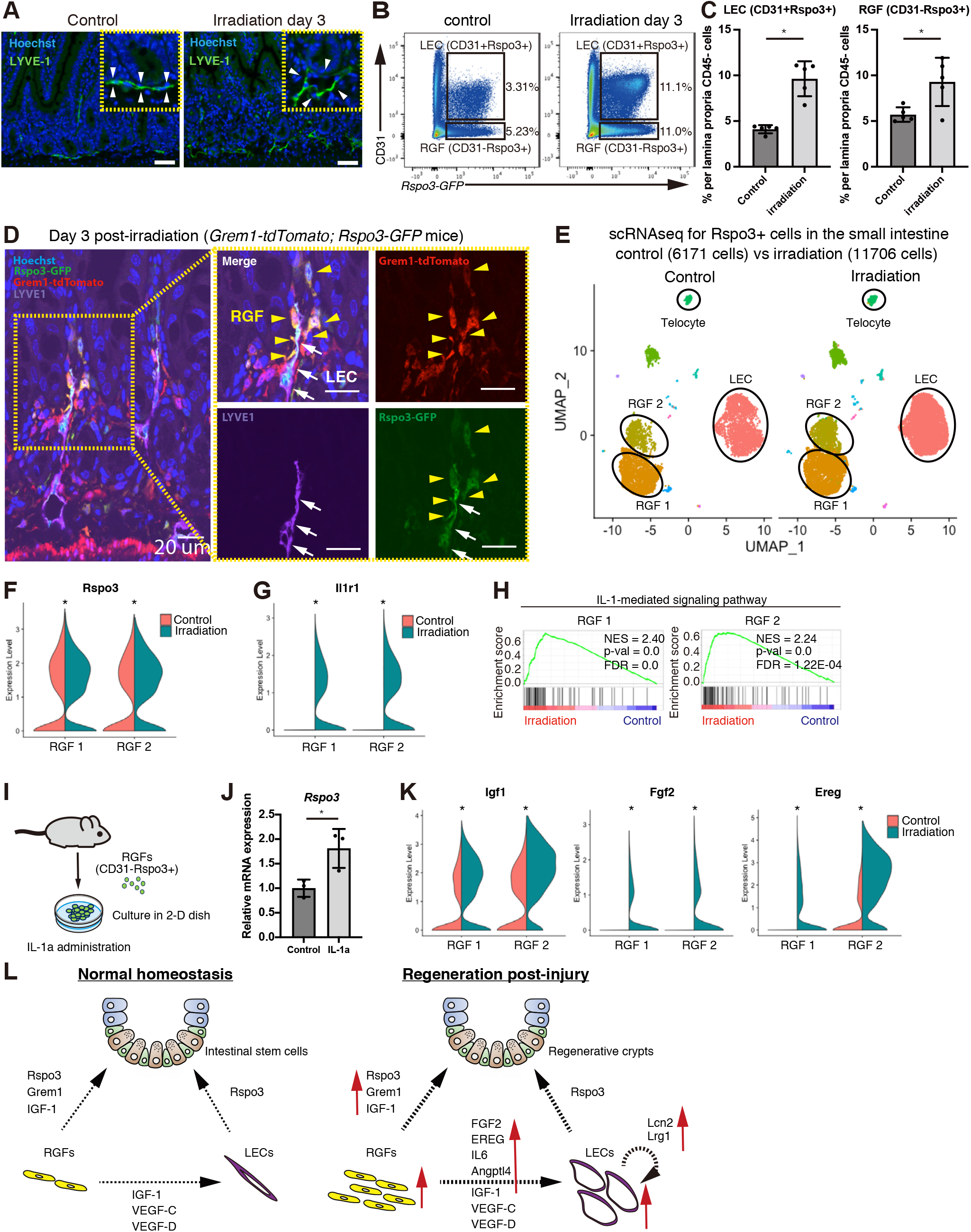
LECs and RGFs expand to facilitate epithelial regeneration after irradiation induced damage. (**A)** Immunofluorescence (IF) for LYVE1 in the small intestine post-irradiation. The images represent one of 6 biological replicates per group. **(B-C)** Flow cytometry (**B**) and quantification (**C**) of CD31+Rspo3-GFP+ LECs and CD31-Rspo3-GFP+ RGFs from the small intestinal EpCAM^−^CD45^−^ stromal cells. n = 5 mice per group. **(D)**Confocal microscopy of IF for Rspo3-GFP, Grem1-tdTomato, and LYVE1 in the small intestine of *Rspo3-GFP; Grem1-tdTomato* mouse day 3 post-irradiation. Yellow arrowheads indicate expanded RGFs. White arrows indicate LECs. The images represent one of 3 biological replicates. **(E)** UMAP of scRNA-seq of sorted small intestinal Rspo3-GFP+ cells 3 days post-irradiation (Control, n = 2 mice and 6,171 cells; irradiation, n = 2 mice and 11,706 cells; multi-dataset integration methods). RGF, RGFs. **(F)** Violin plots for Rspo3 expression. **(G)** Violin plots for Il1r1 expression. **(H)** GSEA of IL-1-mediated signaling pathway genes. **(I)** Schematic of IL-1a administration to RGFs. **(J)** qRT-PCR of Rspo3 mRNA expression from RGFs after IL-1a administration. n = 3 mice per group. **(K)** Violin plots for Igf1, Fgf2, and Ereg expression. **(L)** Model of how LECs and RGFs support ISCs in homeostasis and injury. RGF 1 control, 2,226 cells from 2 mice; RGF 1 irradiation 3,471 cells from 2 mice; RGF 2 control, 812 cells from 2 mice; RGF 2 irradiation, 1,380 cells from 2 mice (**F**, **G**, **K**). Unpaired two-tailed t-tests (**C**, **J**). Wilcox test (**F**, **G**, **K**). Data are mean ± SD. *p < 0.05. Scale bar, 50 μm (**A**) and 20 μm (**D**).

To gain molecular insights into how Rspo3+ stromal cells expand during regeneration and influence ISCs, we performed scRNA-seq in sorted *Rspo3-GFP*+ cells three days post-irradiation. *Rspo3-GFP*+ stromal cells from the irradiation mice clustered separately from control mice, reflecting injury-induced changes (Figure S6A). We then leveraged multi-dataset integration (Stuart et al., 2019) to detect corresponding clusters from the normal and irradiated samples. Similar to what we observed in the unperturbed intestine (Figure 2A), major sources of the Rspo3+ cells after irradiation include LECs and RGFs, the latter of which consists of RGF 1 (Cd81+Adamdec1−) and RGF 2 (Cd81− Adamdec1+) (Figures 6E and S6B). Irradiation not only increased the RGF numbers (Figures 6B-D), but also significantly upregulated *Rspo3* expression in both RGF subsets (Figure 6F).

We, then, performed differential gene expression and GSEA analysis between irradiated and control mice in each RGF subset. Among the top differentially expressed genes in them was *Il1r*, whose expression while low in homeostasis was notably elevated after irradiation (Figures 6G and 6H). Interesting, recent work has implicated macrophage derived IL-1 as a signal that activates IL-1R1 mediated signaling in mesenchymal cells to boost to Rspo3 production as a mechanism for improving intestinal recovery after dextran sulfate sodium–induced colitis (Cox et al., 2021). To test whether IL-1 signaling pathway triggers RSPO3 production in RGFs, we administered IL-1a to cultured RGFs (Figure 6I) and observed that *Rspo3* expression was significantly activated upon IL-1a exposure (Figure 6J), linking IL-1 mediated signaling to augmented RSPO3 production in RGFs during regeneration.

In addition to *Il1r*, *Igf1*, *Fgf2*, and *Ereg* were among the most highly differentially expressed genes after irradiation in RGFs (Figure 6K). The induction of these growth factors may support the maintenance or expansion of LECs and ISCs as IGF-1 and FGFs are requisite growth factors for propagating LECs *in vitro* (Bjorndahl et al., 2005; Chang et al., 2004; Lokmic, 2016) (Figure S5E) and present in intestinal organoid media (Fujii et al., 2018). GSEA analysis also showed upregulation of gene sets in “HALLMARK_ANGIOGENESIS” (Figure S6C) and the expression of *Il6* and *Angptl4-*–factors that facilitate angiogenesis (Cohen et al., 1996; Gopinathan et al., 2015)—as being upregulated in irradiated RGFs (Figure S6D), raising the possibility that RGFs may help coordinate the response of LECs and ISCs by providing growth/angiogenic factors during regeneration. Similar to Rspo3, *Il6* and *Angptl4* upregulation occurs in an IL1a dependent manner in cultured RGFs (Figures S6E and S6F), indicating the IL1a may elicit many of the adaptive changes noted in RGFs. Lastly, in LECs, *Lcn2* and *Lrg1*, other essential regulators of angiogenesis (Nguyen et al., 2020; Wang et al., 2013; Yang et al., 2013), are among the most differentially upregulated genes after irradiation (Figure S6G), implying additional layers of autocrine mechanisms underlying LEC expansion. Collectively, our data support a model whereby RGFs, partially through IL1a mediated signaling, serve as a hub to coordinate ISC and LEC adaptation and regeneration to injury through the heightened production of RSPO3 and growth/angiogenic factors such as IGF-1, FGF2 and Angptl4 (Figure 6L).

## DISCUSSION

Understanding the cell types and trophic/growth factors that support stem cells in diverse tissues is of paramount importance for delineating how stem cells coordinate tissue remodeling or regeneration with organismal needs. This process is of particular importance in the intestine where rapidly renewing Lgr5+ ISCs dynamically sustain the turnover of the intestine in homeostasis, aging, and diverse dietary inputs (Barker et al., 2007; Beyaz et al., 2016; Cheng et al., 2019; Mana et al., 2021; Pentinmikko et al., 2019; Sato et al., 2009). Here, we propose that two mesenchymal populations, namely LECs and RGFs, that reside in close proximity to ISCs play a critical and previously unappreciated role in as the predominant sources of RSPO3, the ligand of Lgr4/5 receptors on ISCs and early progenitors. By leveraging newly engineered fluorescent Rspo3 and Grem1 reporter mice, we find that RGFs, which include previously characterized CD81^+^Pdgfra^low^ trophocytes, produce both Rspo3 and Grem1 and are in close physical proximity to LECs, the other primary source of Rspo3.

Addressing the necessity of Rspo3 in ISC maintenance with genetically engineered mouse models has been challenging as previous studies have demonstrated that it has at best a moderate impact on intestinal homeostasis and integrity (Greicius et al., 2018; Harnack et al., 2019). Possible explanations include the following: First, Rspo3 has redundant cellular sources, namely LECs and RGFs. To date, none of the earlier experimental models disrupted Rspo3 in both of these populations or specifically in LECs. For instance, while we find that a small subset of telocytes or Pdgfra^high^ subepithelial myofibroblasts (SEMFs) express Rspo3 as was previously described, these populations likely represent minor populations for Rspo3 production in the intestine based on our and others scRNAseq analysis (Figures 2A, 2E, S3A, and S3B). Although Rspo3 loss with a PDGFRA-Cre allele to model deletion in SEMFs decreased ISCs numbers and only impacted intestinal morphology after chemical injury, the PDGFRA promoter has overlapping expression in other mesenchymal population such as RGFs, a key RSPO3 source (Figures 2A, 2E, S3A, and S3B); thus, it is likely that this model evaluated Rspo3 loss in more than one stromal cell population but not in LECs. Second, although Rspo3 is the most potent and highly expressed Rspo in the intestine, other Rspos (e.g., Rspo1, Rspo2) may compensate for chronic, long-term Rspo3 loss, especially when Rspo3 is deleted prenatally with Cre only drivers. Conditional Rspo3 loss in either LECs and RGFs decreased ISC numbers and when co-deleted in both cellular populations led to shortened villi-crypt units, near complete absence of Lgr5+ ISCs, and ulcerated mucosa after irradiation induced damage. These results illustrate that LECs and RGFs represent redundant Rspo3 sources. Lastly, our finding that Rspo3 loss in the ISC niche disrupts intestinal homeostasis is consistent with a recent study using a novel *Lgr5-2A-DTR* mouse model, where Lgr5+ cells ablation compromises intestinal integrity (Tan et al., 2021).

In addition to providing homeostatic support to ISCs, we find that RGFs and LECs help aid ISC mediated repair after irradiation. In response to injury, both populations doubled in size relative to all mucosal cells to increase the Rspo3 niche pool to help bolster ISC function. Furthermore, we find that RGFs in part boost Rspo3 production in a IL-1r mediated manner (Figure 6G-J), a pathway that has been implicated in regeneration and whose ligand, IL-1a, is released after damage in the colon, skin and lung (Cox et al., 2021; Katsura et al., 2019; Lee et al., 2017). Interestingly, injury adapted RGFs and LECs also augment their expression of growth factors (e.g. in RGFs, *Igf1 (Bjorndahl et al., 2005; Fujii et al., 2018; Lokmic, 2016), Fgf2 (Chang et al., 2004; Fujii et al., 2018; Lokmic, 2016)*) and angiogenic factors (in LECs, *Lcn2* (Nguyen et al., 2020; Yang et al., 2013), *Lrg1 (Wang et al., 2013)*; in RGFs, *Il6* (Gopinathan et al., 2015), *Angptl4* (Cohen et al., 1996)), indicating that RGFs and LECs interact in injury-induced repair through juxtacrine and autocrine mechanisms to expand their numbers.

Finally, a recent study showed that hair follicle stem cells remodel lymphatics (Gur-Cohen et al., 2019; Pena-Jimenez et al., 2019); yet, whether LECs can reciprocally influence stemness was not explored. Our data demonstrate that LECs serve as a critical in vivo reservoir for Rspo3 production for ISCs in homeostasis and injury-mediated repair and may plausibly do so in other tissues, especially those maintained by Lgr5 expressing stem cells. While LECs and RGFs play a critical role in sustaining ISCs in homeostasis and injury, future work will be required to ascertain the role that these niche cells play in coordinating stem cell function in response to diverse diet-induced organismal physiologies, aging, and disease states.

## ACKNOWLEDGEMENTS

We thank the Swanson Biotechnology Center at the Koch Institute, which encompasses the Flow Cytometry, Histology, and Genomics & Bioinformatics Core facilities (NCI P30-CA14051). We thank Charlie Whittaker and Dikshant Pradhan for analysis and helpful discussions regarding RNA sequencing data. We thank the Department of Comparative Medicine for mouse husbandry support. We thank Sven Holder and members of the Hope Babette Tang (1983) Histology Facility for substantial histology support. We thank members of the Yilmaz laboratory for discussions, Kerry Kelley for laboratory management, and Liz Galoyan for administrative assistance. N.G. is supported by Postdoctoral Fellowship for Research Abroad from Japan Society for the Promotion of Science. Ö.H.Y. is supported by R01CA211184, R01CA034992, and U54CA224068; a Pew-Stewart Trust scholar award; the Kathy and Curt Marble cancer research award; a Koch Institute-Dana-Farber/Harvard Cancer Center Bridge Project grant; and AFAR. Ö.H.Y. receives support from the MIT Stem Cell Initiative.

## AUTHOR CONTRIBUTION

N.G. conceived, designed, performed, interpreted all of the experiments and wrote the manuscript with Ö.H.Y. S.I. provided experimental support. V.D. provided human histopathological samples with diagnostic information and assisted the interpretation of immunohistochemistry. All of the authors assisted in the interpretation of the experiments and the writing of the paper.

## SUPPLEMENTARY FIGURE LEGENDS

**Figure S1. Generation and validation of *Lgr5-2A-GFP-2A-CreERT2* mice, Related to Figure 1 (A)** Rspo3 mRNA expression in the small intestine and colon by ISH. (**B)** Schematic (top) of new *Lgr5-2A-GFP-2A-CreERT2* mice. Immunofluorescence shows that Lgr5-GFP is expressed in all of the crypts of both the small intestine and colon. The image represents one of 6 biological replicates. **(C)** Lineage tracing using *Lgr5-2A-GFP-2A-CreERT2; Rosa-LSL-tdTomato* mice reveals that *Lgr5-GFP*+ cells self-renew for the long-term and give rise to specialized progeny cells both in the small intestine and colon. The image represents one of 3 biological replicates. Scale bar, 50 μm (**A**, **B**, **C**).

**Figure S2. Rspo3 is expressed by LECs and by a subset of Pdgfra+ fibroblasts that co-express Grem1, Related to Figure 1 and Figure 2 (A)** Uniform manifold approximation and projection (UMAP) of scRNA-seq of small intestinal stromal cells (McCarthy et al., 2020, reanalysis). Rspo3 is expressed by Lyve+ LEC cluster and by a subset of Pdgfra+ fibroblasts cluster (top). Rspo3+ cells in the Pdgfra+ fibroblasts cluster co-express Grem1. **(B)** UMAP of scRNA-seq of colonic stromal cells (Kinchen et al., 2018, reanalysis). Rspo3 is expressed by Lyve+ LEC cluster and by a subset of Pdgfra^low^ fibroblasts (both CD81-Pdgfra^low^ fibroblasts and CD81+Pdgfra^low^ fibroblasts). A subset of Rspo3+ cells in the Pdgfra^low^ fibroblasts clusters co-express Grem1.

**Figure S3. Rspo3+Grem1+ cell distribution in Pdgfra+ fibroblasts, Related to Figure 2 (A)** UMAP of scRNA-seq of small intestinal Pdgfra+ cells (McCarthy et al., 2020, reanalysis). The expressions of Cd81 and Adamdec1 are mutually exclusive (Top). Rspo3+ cells are detected both in the Cd81^+^Adamdec1^−^Pdgfra^low^ trophocyte and Cd81^−^Adamdec1^+^Pdgfra^low^ fibroblast clusters, and these Rspo3+ cells frequently co-express Grem1. **(B)** UMAP of scRNA-seq of colonic Pdgfra+ cells (Brugger et al., 2020, reanalysis). The expressions of Cd81 and Adamdec1 are mutually exclusive (Top). Rspo3+ cells are detected both in the Cd81^+^Adamdec1^−^Pdgfra^low^ trophocyte and CD81^−^Adamdec1^+^Pdgfra^low^ fibroblast clusters, and these Rspo3+ cells frequently co-express Grem1. **(C)** UMAP of scRNA-seq of human colonic stromal cells (Kinchen et al., 2018, reanalysis). The expressions of CD81 and ADAMDEC1 are ubiquitous in PDGFRA^low^ fibroblasts and not mutually exclusive (Top). RSPO3+GREM1+ cells are detected in a subset of PDGFRA^low^ fibroblasts.

**Figure S4. RNA-seq on CD31-Rspo3-GFP-, CD31+Rspo3-GFP-, CD31-Rspo3-GFP+, and CD31+Rspo3-GFP+ cells, Related to Figure 3 (A**) GSEA of CD81^+^Pdgfra^low^ trophocyte markers, Ackr4+ fibroblast markers, and lymphatic markers. FDR, false-discovery rate; NES, normalized enrichment score. (**B**) The heatmap of single sample GSEA (ssGSEA). (**C-I)**The heatmap of gene expressions for angiogenic factors (**C**), growth factors (**D**), BMPi (**E**), BMP (**F**), Rspo (**G**), Wnt antagonist (**H**), and Wnt (**I**) using RNA-seq on CD31-Rspo3-GFP-, CD31+Rspo3-GFP-, CD31-Rspo3-GFP+, and CD31+Rspo3-GFP+ cells from the small intestine and colon. n = 3 mice per group.

**Figure S5. Expression pattern of Grem1 and Rspo3 and establishment of Rspo3+ stromal cell culture, Related to Figure 3 and Figure 4 (A)** Schematic of *Grem1-tdTomato-CreERT2* mouse. **(B)** Confocal microscopy of immunofluorescence (IF) for Grem1-tdTomato and Desmin, showing that Grem1-tdTomato is expressed by a subset of lamina propria and submucosal stromal cells, but is also strongly expressed by Desmin+ muscularis propria and muscularis mucosa cells. **(C)** IF for Rspo3-GFP, Grem1-tdTomato, and Adamdec1 reveals that RGFs close to the crypt bottoms do not express Adamdec1 whereas RGFs infrequently detected in the villous core next to lacteals co-express Adamdec1. (**D**) Grem1+Rspo3− cells at the middle-top zone of colonic crypts. lower magnification image is adapted from Figure 3E. **(E-F)** 2-D (**E**) and 3-D (**F**) culture of sorted *Rspo3-GFP*+ stromal cells from the small intestine. **(G-H)** Images (**G**) and quantification (**H**) of 2-D culture of sorted CD31^−^*Rspo3-GFP*^−^, CD31^+^*Rspo3-GFP*^−^, CD31^+^*Rspo3-GFP*^+^, and CD31^−^*Rspo3-GFP*^+^ cells from the small intestine at day 5. RGF, RGFs. (**I**) Schematic of heterotypic co-culture of Rspo3+ stromal cells and the intestinal epithelial crypts. (**J-K**) Representative images (**J**) and quantification (**K**) of co-culture of small intestinal Rspo3+ stromal cells (0, 2×10^4^, 1×10^5^) and the crypts in the culture media supplemented with Noggin, but not with Rspo. n = 12 - 13 from 3 mice per group. Arrows indicate organoid formation. One-way analysis of variance (ANOVA) with post-hoc Tukey’s multiple comparison (**H, K)**. Data are mean ± SD. *p < 0.05. Scale bar, 20 μm (**B**, **C, E**, **F**, **G**, **J**), 50 μm (**D**).

**Figure S6. LECs and RGFs under normal homeostasis and regeneration after injury, Related to Figure 6 (A)** UMAP of scRNA-seq of small intestinal *Rspo3-GFP*+ cells 3 days post-irradiation before (left) and after (right) applying multi-dataset integration methods (Control, n = 2 mice and 6,171 cells; irradiation, n = 2 mice and 11,706 cells). (**B**) Relative expression of *Cd81* and *Adamdec1* on *the* UMAP plot of Rspo3+ cells from both control and irradiation mice. n = 17,877 cells from 4 mice. **C**. GSEA of HALLMARK_Angiogenesis genes in irradiation vs control mice from RGF 1 cluster (left) and RGF 2 cluster (right) of scRNA-seq. (**D**) Violin plots for *Il6* and *Angptl4*. (**E**) Schematic of the experiments of IL-1a administration to RGF culture. Adapted from Figure 5I. (**F**) qRT-PCR of *Il6* and *Angptl4* mRNA expression from RGFs 24 hours after IL-1a administration. n = 3 mice per group. (**G**) Violin plots for *Lcn2* and *Lrg1* expression comparing control and irradiation mice in LEC cluster of scRNA-seq. Unpaired two-tailed t-tests (**F**). Wilcox test (**D**, **G**). Data are mean ± SD. *p < 0.05.

**Supplementary Video 1: 3-D reconstitution of confocal microscopy images of immunofluorescence for Lgr5-GFP and LYVE1 in the mouse small intestine**

**Supplementary Table 1: Custom gene sets for Gene Set Enrichment Analysis (GSEA)**

## Methods

### RESOURCE AVAILABILITY

#### Lead contact

Further information and requests for resources and reagents should be directed to and will be fulfilled by Ömer H. Yilmaz (, (Ö.H.Y.)).

#### Materials availability

All in-house generated mouse strains generated for this study are available from the Lead Contacts with a completed Materials Transfer Agreement.

#### Data and code availability

All datasets generated in this study will be made available on GEO upon publication. All relevant data supporting the findings of this study are also available from Ömer H. Yilmaz or Norihiro Goto upon request.

### EXPERIMENTAL MODEL AND SUBJECT DETAILS

#### Animal models

Mice were under the husbandry care of the Department of Comparative Medicine in the Koch Institute for Integrative Cancer Research. All procedures were conducted in accordance with the American Association for Accreditation of Laboratory Animal Care and approved by MIT’s Committee on Animal Care. *Lgr5-2A-EGFP-2A-CreERT2* mice was generated by inserting P2A-EGFP-T2A-CreERT2 cassette in the endogenous Lgr5 gene locus immediately at the 5′ end of the endogenous stop codon using CRISPR-Cas9 technology and zygote microinjection. *Grem1-2A-tdTomato-2A-CreERT2* mouse was generated by inserting P2A-tdTomato-T2A-CreERT2 cassette in the endogenous *Grem1* gene locus immediately at the 5′ end of the endogenous stop codon using CRISPR-Cas9 technology and zygote microinjection. Successful targeting was validated by Southern blotting and PCR analysis. *Rspo3-GFP* BAC transgenic mouse was obtained from the Gene Expression Nervous System Atlas Project (Heintz, 2004) and backcrossed to C57BL/6J mice for more than 10 generations. *Rspo3 flox* mice (Neufeld et al., 2012) (JAX strain 027313), *Prox1-CreERT2* mice (Srinivasan et al., 2007) (JAX strain 022075), and *Rosa26-LSL-tdTomato* mice (JAX strain 007914) were obtained from the Jackson Laboratory. These mice were maintained in a C57BL/6 background. In this study, both male and female mice were used at the ages of 8-12 wks. For induction of Cre-mediated recombination, 200 μl of 20 mg/ml tamoxifen in corn oil (Figure S1C), or 100 μl of 20 mg/ml tamoxifen over 4 consecutive days (Figures 5A-M) were intraperitoneally injected.

#### Human intestinal samples

Human duodenal biopsies that were diagnosed as normal were obtained from 10 patients. The Massachusetts General Hospital (MGH) Institutional Review Board committee approved the study protocol.

### METHOD DETAILS

#### Immunohistochemistry (IHC) and immunofluorescence (IF)

Tissues were fixed in 10% formalin, paraffin embedded and sectioned in 4-5 micron sections as previously described (Beyaz et al., 2016; Cheng et al., 2019; Mana et al., 2021). Antigen retrieval was performed using Borg Decloaker RTU solution (Biocare Medical, BD1000G1) and a pressurized Decloaking Chamber (Biocare Medical, NxGen). Antibodies and respective dilutions used for immunohistochemistry are as follows: rabbit monoclonal anti-OLFM4 (1:10,000, CST, 39141) and goat polyclonal anti-GFP (1:500, abcam, ab6673). Biotin-conjugated secondary donkey anti-rabbit or anti-goat antibodies were used (1:500, Jackson ImmunoResearch). Vectastain Elite ABC immunoperoxidase detection kit (Vector Laboratories, PK6100) was followed by Signalstain DAB substrate kit for visualization (CST, 8049S). All antibody dilutions were performed in Signalstain Antibody Diluent (CST, 8112L). The following primary antibodies were used for immunofluorescence: goat polyclonal anti-GFP (1:500, abcam, ab6673), rat monoclonal anti-LYVE1 (1:1000, Thermo Fisher Scientific, 14-0443-82), rabbit polyclonal anti-LYVE1 (1:1000, Sigma Aldrich, HPA042953), mouse monoclonal anti-Desmin (1:200, abcam, ab6322), mouse monoclonal anti-ADAMDEC1 (1:100, Thermo Fisher Scientific, 6C4), and rabbit polyclonal anti-RFP (1:400, Rockland 600-401-379). Alexa Fluor secondary antibodies, anti-goat 488, anti-mouse 488, anti-rabbit 568, anti-mouse 647, and anti-rat 647 (1:500, Thermo Fisher Scientific), were used for visualization. Slides were stained with Hoechst for 10 min and covered with Prolong Gold (Life Technologies, P36930) mounting media. Images were acquired using a Nikon Eclipse 90i upright microscope equipped with a Hamamatsu Orca-ER CCD camera, and APC line 1200 light source. For the acquisition of high-resolution confocal images, Nikon A1R Ultra-Fast Spectral Scanning Confocal Microscope was used.

#### Intestinal stromal cell isolation and flow cytometry

Intestinal stromal cells were isolated as previously reported with a slight modification (McCarthy et al., 2020b; Ogasawara et al., 2018). The small intestines and the colons were removed, washed with cold PBS, opened longitudinally, cut into approximately 5 mm pieces, and then incubated with PBS plus EDTA (10 mM) for 40 min at 37°C. Crypts were mechanically removed from the tissue by shaking and washing with PBS. The remaining tissue was digested on a shaker at 400 rpm for 40 min at 37°C in 100 μg/ml Liberase TM (Roche, 5401127001) and 100 μg/ml DNAse I (Roche, 10104159001) diluted in RPMI 1640 media (Thermo Fisher Scientific, 11875093), containing 2% fetal bovine serum (FBS). Every 20 min, the tissue suspension was passed through an 14-gauge needle using a 10-ml syringe for mechanical dissociation. Extracted cells were passed through a 40 μm filter, centrifuged at 500 g for 5 min, and washed with RPMI 1640 medium containing 2% FBS. Cells were resuspended in ACK lysis buffer (Thermo Fisher Scientific, A1049201), incubated on ice for 4 minutes to remove red blood cells, washed with RPMI-1640 medium containing 2% FBS, and resuspended in the same buffer. Dissociated single cells were treated with the following antibody cocktail for flow cytometry analysis: CD45-PE (1:200, ThermoFisher,12-0451-83), CD31-eFluor450 (1:500, ThermoFisher, 48-0311-82), EpCAM-APC (1:300, ThermoFisher, 17-5791-82). 7AAD (ThermoFisher, A1310) was used a viability dye to exclude dead cells from the analysis. Fluorescence-activated cell sorting was performed using a FACS Aria II (BD Biosciences). The data were analyzed using FlowJo software (version 10, TreeStar) and FACSDiva software (version 8.0, BD Biosciences).

#### Culture of Rspo3+ stromal cells

Sorted Rspo3+, CD31-Rspo3-, CD31+Rspo3-, CD31-Rspo3+, and CD31+Rspo3+ cells were centrifuged at 500 g for 5 min and resuspended in 500 μl endothelial cell media: EGM-2 MV Bullet Kit (Lonza, cc-3202) supplemented with 50 ng/mL VEGF-C (R&D, 2179-VC-025). The kit contains EGF, hydrocortisone, gentamicin (GA-1000), FBS, VEGF, FGF-b, IGF-1, and ascorbic acid. Cells were plated in a fibronectin-coated 24-well plate in total volume of 500 μl with 2×10^4^ cells per well. Endothelial cell media were changed every other day. Once the cells reached 80% confluence, the cells are detached from the wells with Accutase (STEMCELL technologies, 07920) for passage and co-culture with the intestinal crypts. For IL-1a administration experiments, sorted CD31-Rspo3+ cells were cultured and recombinant murine IL-1a (10 ng/ml, Peprotech, 211-11A) was administered to each well 24 hours prior to RNA extraction.

#### Intestinal crypt isolation and co-culture of intestinal crypts with stromal cells

The small intestines were removed, washed with cold PBS, opened longitudinally and then incubated on ice with PBS plus EDTA (10 mM) for 45 min. Tissues were then moved to PBS. Crypts were then mechanically separated from the connective tissue by shaking, and then filtered through a 100 μm mesh into a 50 mL conical tube to remove villus material and tissue fragments. For co-cultures, 100 crypts were cultured in 48-well tissue culture plates loaded with 10 μl drops of Matrigel (Corning) together with 1×10^5^ stromal cells detached from the short-term culture after sorting as described above. Endothelial cell media supplemented with B27 1X (Life Technologies) and Y-27632 dihydrochloride monohydrate 10 μM (Sigma-Aldrich) were added to the wells with or without 100 ng/mL Noggin (Peprotech, 250-38). Culture media were changed every other day, and cell cultures were maintained at 37°C in fully humidified chambers containing 5% CO_2_.

#### qRT-PCR and in situ hybridization

Total RNA was isolated using RNeasy Mini Kit (QIAGEN, 74104) or RNeasy Micro Kit (QIAGEN, 74004) according to the manufacturer’s instructions. RNA was converted to cDNA using qScript cDNA SuperMix (Quantabio, 95048-100). Quantitative RT-PCR (qRT-PCR) reaction was performed using cDNA with SYBR green fast mix (Quantabio, PerfeCTa, 95072-012) on a Roche lightcycler (Roche, LightCycler 480 II). The following primers used for qRT-PCR: *Gapdh* forward, 5’-AGGTCGGTGTGAACGGATTTG-3’; *Gapdh* reverse, 5’-TGTAGACCATGTAGTTGAGGTCA-3’; *Rspo3* forward, 5’-ATGCACTTGCGACTGATTTCT-3’; *Rspo3* reverse, 5’-GCAGCCTTGACTGACATTAGGAT-3’; *Il6* forward, 5’-TAGTCCTTCCTACCCCAATTTCC-3’; *Il6* reverse, 5’-TTGGTCCTTAGCCACTCCTTC-3’; *Agptl4* forward, 5′-CATCCTGGGACGAGATGAACT-3′; *Agptl4* reverse, 5′-TGACAAGCGTTACCACAGGC-3′.

Single molecule in situ hybridization was performed using Advanced Cell Diagnostics RNAscope 2.5 HD Detection Kit. The in situ hybridization probes used in this study are as follows: Mm-Rspo3 (Ref 402011), Mm-Lgr5 (Ref 312171).

#### RNA-seq data processing and differential expression analysis

Single-end RNA-seq data was used to quantify transcripts from the mm10 mouse assembly with the Ensembl version 98 annotation using Salmon version 1.1.0 (Patro et al., 2017). Paired-end RNA-seq data was used to quantify transcripts from the mm10 mouse assembly with the Ensembl version 101 annotation using Salmon version 1.3.0 (Patro et al., 2017). Gene level summaries were prepared using tximport version 1.21.1, 1.16.1 (Soneson et al., 2015) running under R version 3.6.1, 4.0.2 (R Core Team 2021). Differential expression analysis was done with DESeq2 version 1.24.0 (Love et al., 2014) and differentially expressed genes were defined as those having an absolute apeglm (Zhu et al., 2019) log2 fold change greater than 1 and an adjusted p-value less than 0.05. Data parsing and some visualizations was done using Tibco Spotfire Analyst 7.6.1. Mouse genes were mapped to human orthologs using Mouse Genome Informatics (http://www.informatics.jax.org/) orthology report and preranked Gene Set Enrichment Analysis (Mootha et al., 2003) was done using javaGSEA version 4.1.0 with a custom gene sets or sets from MSigDB version 7.2 (Subramanian et al., 2005).

#### scRNA-seq data processing and analysis

scRNA-seq data was produced using the 10x genomics platform. Data were processed using cell ranger version 6.0.1 with alignment to a modified version of the GENCODE mouse genome (GRCm38), version M23 (Ensembl 98) target provided by 10x genomics. The modification was the addition of the GFP selectable marker. Cellranger filtered data was imported into R version 4.1.0 (R Core Team 2021) and analyzed with Seurat version 4.0.3 (Stuart et al., 2019).

#### Irradiation experiments

Mice were challenged by 12 Gy of irradiation. Tissue was collected 72 hours post-irradiation. Numbers of surviving crypts were enumerated in the proximal jejunum from Hematoxylin and Eosin stained tissue and identified as robust crypt structures with dense nuclei and presence of Paneth cells.

### QUANTIFICATION AND STATISTICAL ANALYSIS

Unless otherwise specified in the figure legends or Method Details, all experiments reported in this study were repeated at least three independent times. For organoid co-culture assays, 2-5 wells per group with at least 3 different mice were analyzed. All sample number (n) of biological replicates and technical replicates, definition of center, and dispersion and precision measures can be found in the figure legends. All values are presented as mean ± SD unless otherwise stated. Intergroup comparisons were performed using unpaired two-tailed t-tests or one-way analysis of variance (ANOVA) with post-hoc Tukey’s multiple comparison. P values of < 0.05 were considered to be significant. Statistical analysis was performed by GraphPad Prism. No sample or animals were excluded from analysis. Age- and sex-matched mice were randomly assigned to groups. Studies were not conducted blind with the exception of all histological analyses. Please note that statistical details are found in the figure legends.

