## Supplementary Figures for "Lymphatics constitute a novel component of the intestinal stem cell niche"

### Figure S1

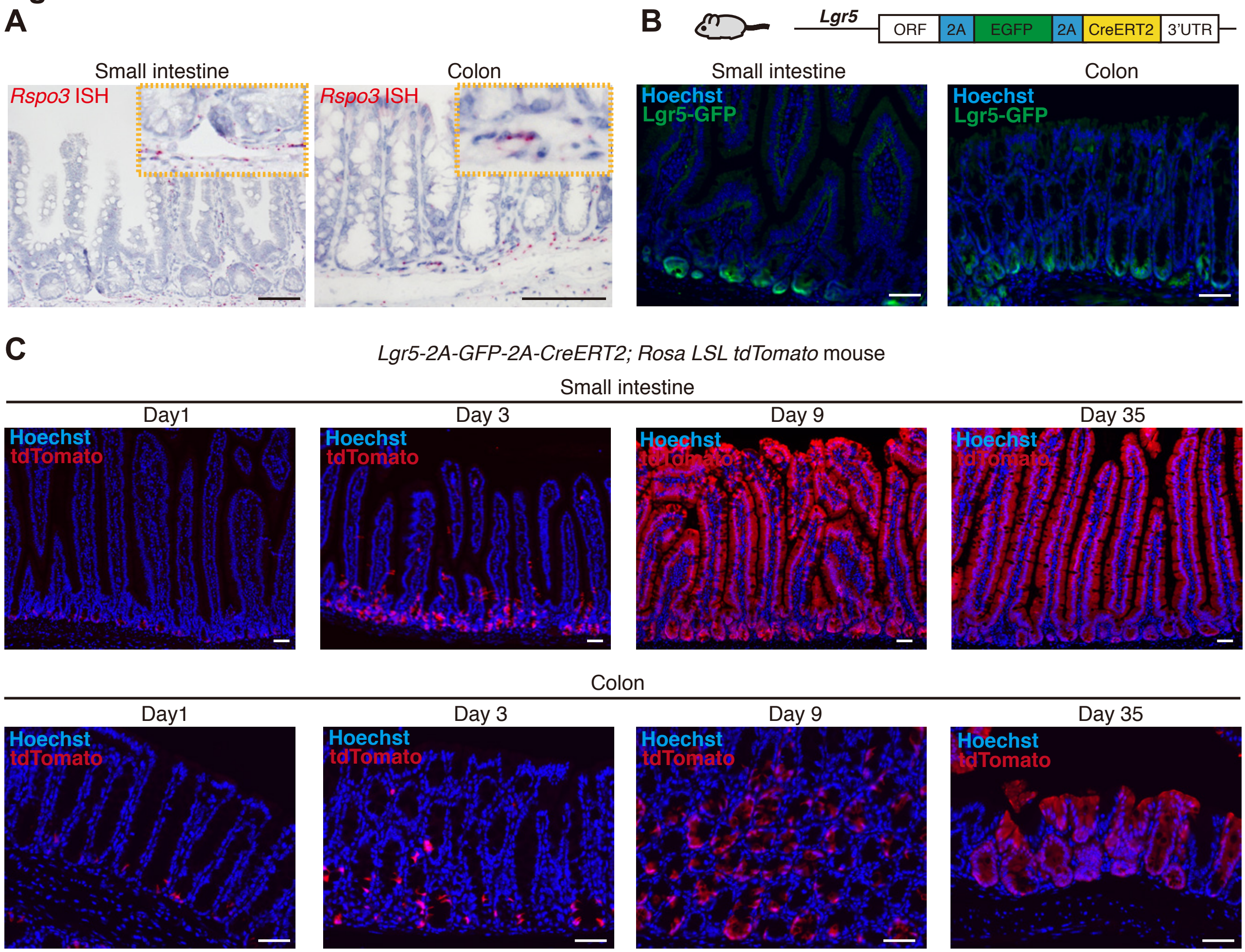

A
Small intestine
Colon
Rspo3 ISH
Rspo3 ISH
B
Lgr5
ORF
2A
EGFP
2A
CreERT2
3'UTR
Small intestine
Colon
Hoechst
Lgr5-GFP
Hoechst
Lgr5-GFP
C
Lgr5-2A-GFP-2A-CreERT2; Rosa LSL tdTomato mouse
Small intestine
Day1
Day 3
Day 9
Day 35
Hoechst
tdTomato
Colon
Day1
Day 3
Day 9
Day 35
Hoechst
tdTomato

### Figure S2

**A** Stromal cells in the mouse small intestine (McCarthy et al., 2020, reanalysis)

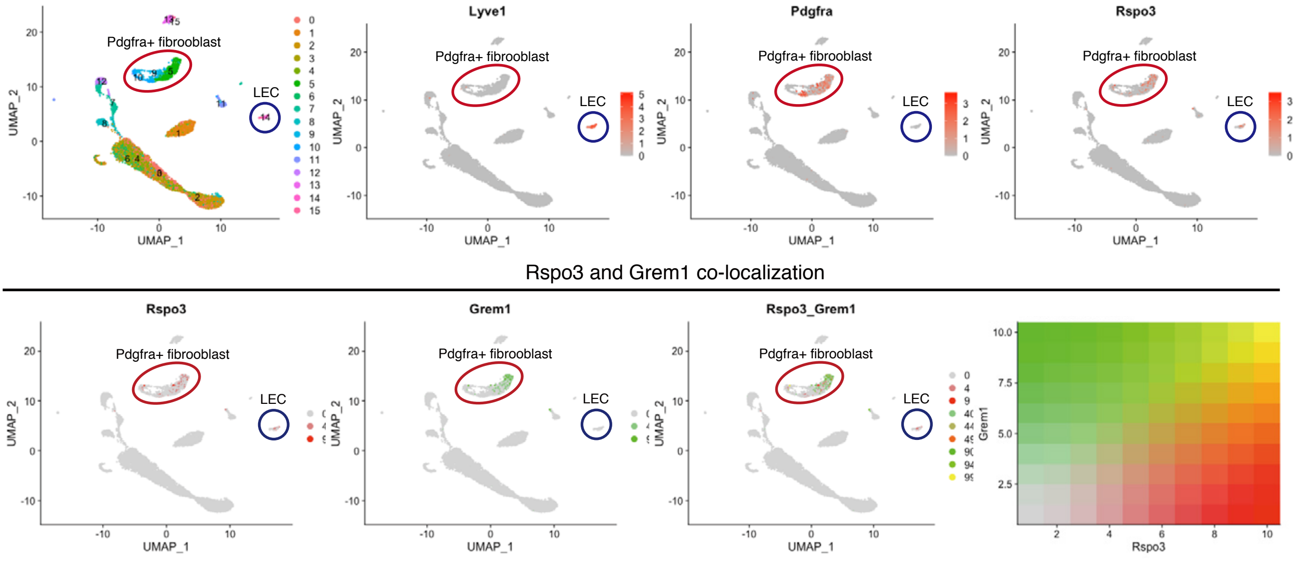


**B** Stromal cells in the mouse colon (Kinchen et al., 2018, reanalysis)

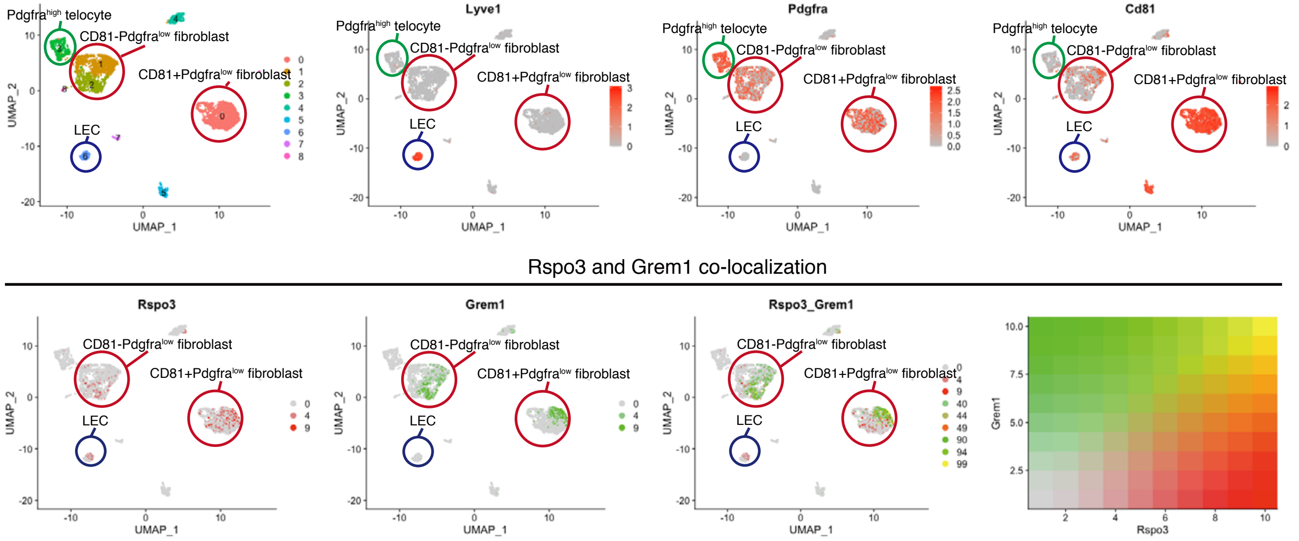

### Figure S3

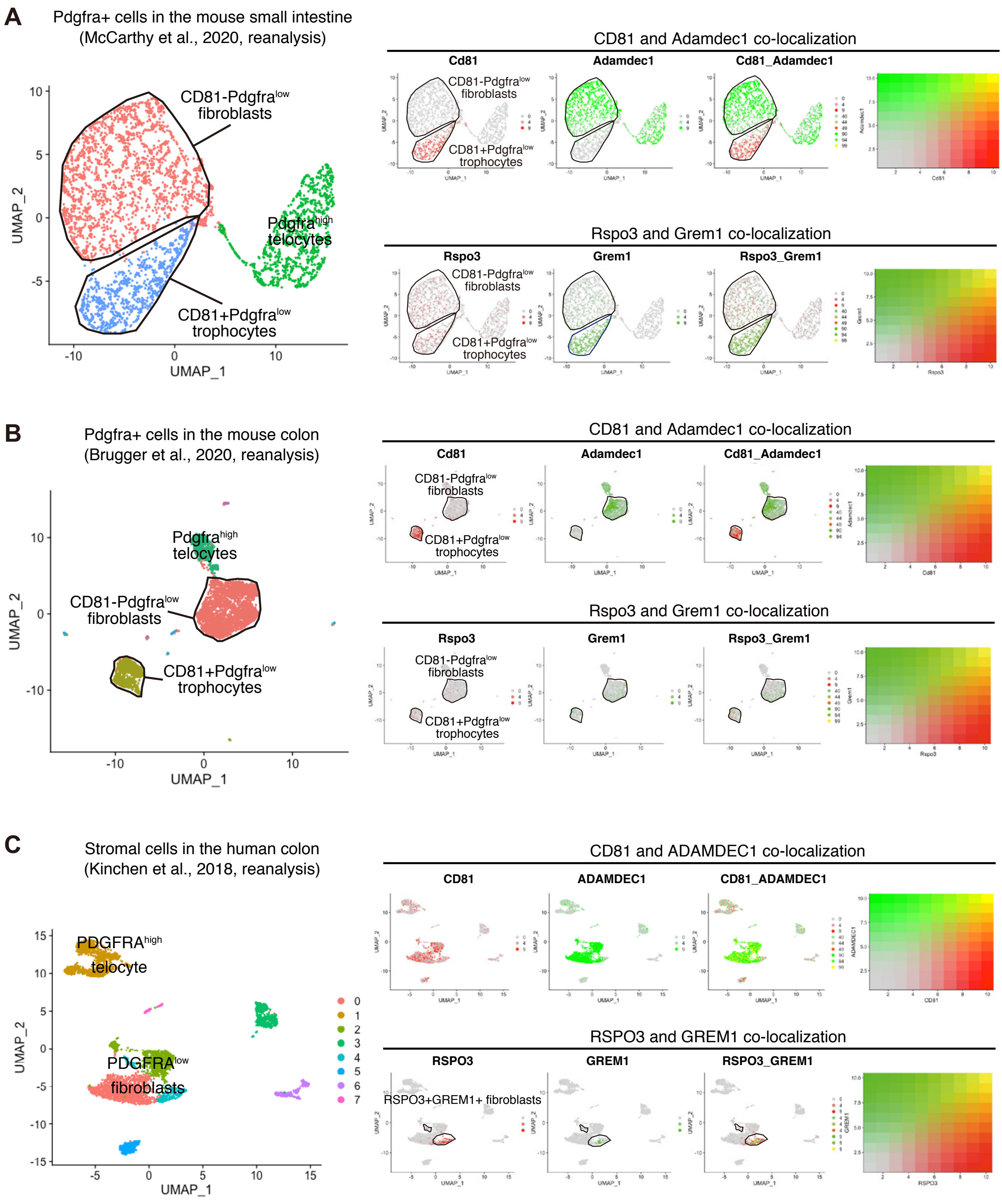
A
Pdgfra+ cells in the mouse small intestine
(McCarthy et al., 2020, reanalysis)
CD81-Pdgfralow fibroblasts
Pdgfrahigh telocytes
CD81+Pdgfralow trophocytes
UMAP_1
UMAP_2
CD81 and Adamdec1 co-localization
Cd81
Adamdec1
Cd81_Adamdec1
Rspo3 and Grem1 co-localization
Rspo3
Grem1
Rspo3_Grem1
B
Pdgfra+ cells in the mouse colon
(Brugger et al., 2020, reanalysis)
Pdgfrahigh telocytes
CD81-Pdgfralow fibroblasts
CD81+Pdgfralow trophocytes
UMAP_1
UMAP_2
CD81 and Adamdec1 co-localization
Cd81
Adamdec1
Cd81_Adamdec1
Rspo3 and Grem1 co-localization
Rspo3
Grem1
Rspo3_Grem1
C
Stromal cells in the human colon
(Kinchen et al., 2018, reanalysis)
PDGFRAhigh telocyte
PDGFRAlow fibroblasts
UMAP_1
UMAP_2
CD81 and ADAMDEC1 co-localization
CD81
ADAMDEC1
CD81_ADAMDEC1
RSPO3 and GREM1 co-localization
RSPO3
GREM1
RSPO3_GREM1
RSPO3+GREM1+ fibroblasts

### Figure S4

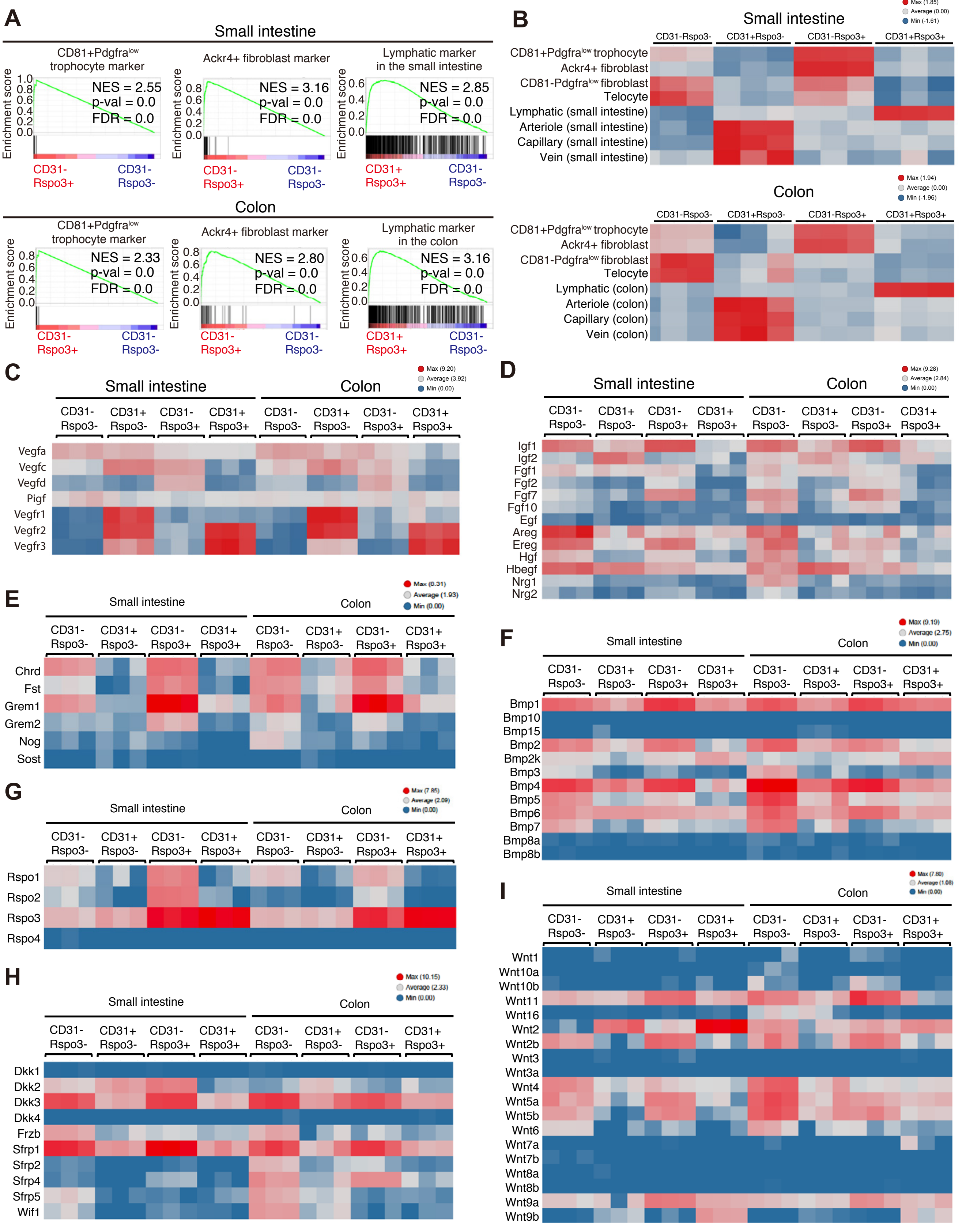
A
Small intestine
CD81+Pdgfra^low trophocyte marker
Ackr4+ fibroblast marker
Lymphatic marker in the small intestine
Enrichment score
NES = 2.55
p-val = 0.0
FDR = 0.0
NES = 3.16
p-val = 0.0
FDR = 0.0
NES = 2.85
p-val = 0.0
FDR = 0.0
CD31- Rspo3+
CD31- Rspo3-
CD31+ Rspo3+
CD31- Rspo3-
Colon
CD81+Pdgfra^low trophocyte marker
Ackr4+ fibroblast marker
Lymphatic marker in the colon
NES = 2.33
p-val = 0.0
FDR = 0.0
NES = 2.80
p-val = 0.0
FDR = 0.0
NES = 3.16
p-val = 0.0
FDR = 0.0
B
Small intestine
Max (1.85)
Average (0.00)
Min (-1.61)
CD31-Rspo3-
CD31+Rspo3-
CD31-Rspo3+
CD31+Rspo3+
CD81+Pdgfra^low trophocyte
Ackr4+ fibroblast
CD81-Pdgfra^low fibroblast
Telocyte
Lymphatic (small intestine)
Arteriole (small intestine)
Capillary (small intestine)
Vein (small intestine)
Colon
Max (1.94)
Average (0.00)
Min (-1.96)
Lymphatic (colon)
Arteriole (colon)
Capillary (colon)
Vein (colon)
C
Small intestine
Colon
Max (9.20)
Average (3.92)
Min (0.00)
CD31- Rspo3-
CD31+ Rspo3-
CD31- Rspo3+
CD31+ Rspo3+
Vegfa
Vegfc
Vegfd
Pigf
Vegfr1
Vegfr2
Vegfr3
D
Max (9.28)
Average (2.84)
Min (0.00)
Igf1
Igf2
Fgf1
Fgf2
Fgf7
Fgf10
Egf
Areg
Ereg
Hgf
Hbegf
Nrg1
Nrg2
E
Max (0.31)
Average (1.93)
Min (0.00)
Chrd
Fst
Grem1
Grem2
Nog
Sost
F
Max (9.19)
Average (2.75)
Min (0.00)
Bmp1
Bmp10
Bmp15
Bmp2
Bmp2k
Bmp3
Bmp4
Bmp5
Bmp6
Bmp7
Bmp8a
Bmp8b
G
Max (7.85)
Average (2.09)
Min (0.00)
Rspo1
Rspo2
Rspo3
Rspo4
H
Max (10.15)
Average (2.33)
Min (0.00)
Dkk1
Dkk2
Dkk3
Dkk4
Frzb
Sfrp1
Sfrp2
Sfrp4
Sfrp5
Wif1
I
Max (7.80)
Average (1.08)
Min (0.00)
Wnt1
Wnt10a
Wnt10b
Wnt11
Wnt16
Wnt2
Wnt2b
Wnt3
Wnt3a
Wnt4
Wnt5a
Wnt5b
Wnt6
Wnt7a
Wnt7b
Wnt8a
Wnt8b
Wnt9a
Wnt9b

### Figure S5

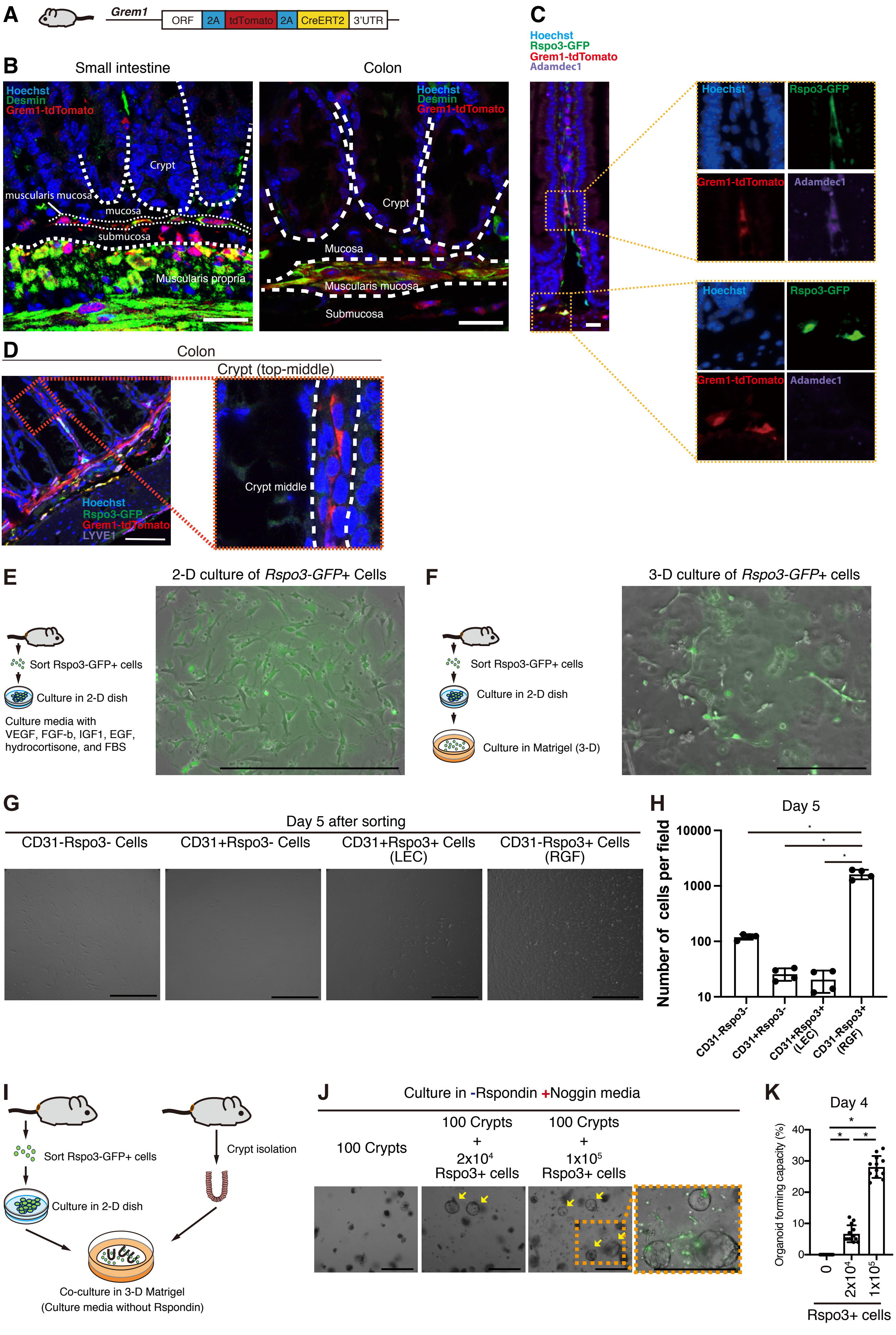
A
Grem1
ORF
2A
tdTomato
2A
CreERT2
3'UTR
B
Small intestine
Hoechst
Desmin
Grem1-tdTomato
Crypt
muscularis mucosa
mucosa
submucosa
Muscularis propria
Colon
Hoechst
Desmin
Grem1-tdTomato
Crypt
Mucosa
Muscularis mucosa
Submucosa
C
Hoechst
Rspo3-GFP
Grem1-tdTomato
Adamdec1
Hoechst
Rspo3-GFP
Grem1-tdTomato
Adamdec1
Hoechst
Rspo3-GFP
Grem1-tdTomato
Adamdec1
D
Colon
Crypt (top-middle)
Hoechst
Rspo3-GFP
Grem1-tdTomato
LYVE1
Crypt middle
E
2-D culture of Rspo3-GFP+ Cells
Sort Rspo3-GFP+ cells
Culture in 2-D dish
Culture media with VEGF, FGF-b, IGF1, EGF, hydrocortisone, and FBS
F
3-D culture of Rspo3-GFP+ cells
Sort Rspo3-GFP+ cells
Culture in 2-D dish
Culture in Matrigel (3-D)
G
Day 5 after sorting
CD31-Rspo3- Cells
CD31+Rspo3- Cells
CD31+Rspo3+ Cells (LEC)
CD31-Rspo3+ Cells (RGF)
H
Day 5
Number of cells per field
10000
1000
100
10
*
*
*
CD31-Rspo3-
CD31+Rspo3-
CD31+Rspo3+ (LEC)
CD31-Rspo3+ (RGF)
I
Sort Rspo3-GFP+ cells
Culture in 2-D dish
Crypt isolation
Co-culture in 3-D Matrigel
(Culture media without Rspondin)
J
Culture in -Rspondin +Noggin media
100 Crypts
100 Crypts + 2x10^4 Rspo3+ cells
100 Crypts + 1x10^5 Rspo3+ cells
K
Day 4
Organoid forming capacity (%)
40
30
20
10
0
*
*
*
0
2x10^4
1x10^5
Rspo3+ cells

### Figure S6

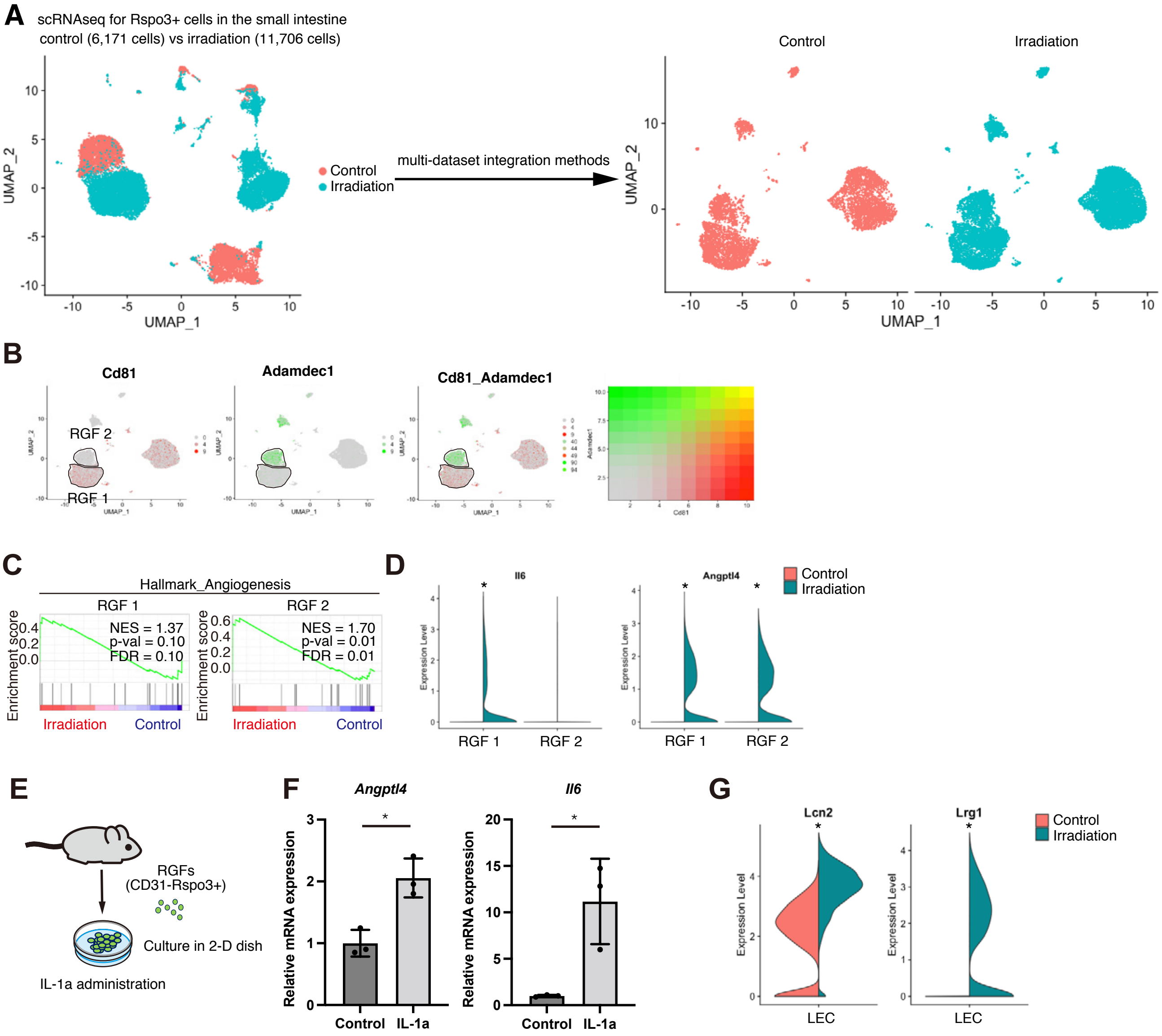
A
scRNAseq for Rspo3+ cells in the small intestine
control (6,171 cells) vs irradiation (11,706 cells)
Control
Irradiation
multi-dataset integration methods
UMAP_1
UMAP_2
B
Cd81
Adamdec1
Cd81_Adamdec1
RGF 2
RGF 1
C
Hallmark_Angiogenesis
RGF 1
RGF 2
NES = 1.37
p-val = 0.10
FDR = 0.10
NES = 1.70
p-val = 0.01
FDR = 0.01
Enrichment score
Irradiation
Control
D
Il6
Angptl4
Expression Level
RGF 1
RGF 2
E
RGFs
(CD31-Rspo3+)
Culture in 2-D dish
IL-1a administration
F
Angptl4
Il6
Relative mRNA expression
Control
IL-1a
G
Lcn2
Lrg1
LEC
